# FBXL4 suppresses mitophagy by restricting the accumulation of NIX and BNIP3 mitophagy receptors

**DOI:** 10.1101/2022.10.12.511867

**Authors:** Giang Thanh Nguyen-Dien, Keri-Lyn Kozul, Yi Cui, Brendan Townsend, Prajakta Gosavi Kulkarni, Soo Siang Ooi, Antonio Marzio, Nissa Carrodus, Steven Zuryn, Michele Pagano, Robert G. Parton, Michael Lazarou, Sean Millard, Robert W. Taylor, Brett M. Collins, Mathew J.K. Jones, Julia K. Pagan

## Abstract

Cells selectively remove damaged or excessive mitochondria through mitophagy, a specialized form of autophagy, to maintain mitochondrial quality and quantity. Mitophagy is induced in response to diverse conditions, including hypoxia, cellular differentiation, and mitochondrial damage. However, the mechanisms by which cells remove specific dysfunctional mitochondria under steady-state conditions to fine-tune mitochondrial content are not well understood. Here, we report that SCF^FBXL4^, an SKP1/CUL1/F-box protein ubiquitin ligase complex, localizes to the mitochondrial outer membrane in unstressed cells and mediates the constitutive ubiquitylation and degradation of the mitophagy receptors NIX and BNIP3 to suppress basal levels of mitophagy. We demonstrate that, unlike wild-type FBXL4, pathogenic variants of FBXL4 that cause encephalopathic mtDNA depletion syndrome (MTDPS13), do not efficiently interact with the core SCF ubiquitin ligase machinery or mediate the degradation of NIX and BNIP3. Thus, we reveal a molecular mechanism that actively suppresses mitophagy via preventing NIX and BNIP3 accumulation and propose that excessive basal mitophagy in the FBXL4-associated mtDNA depletion syndrome is caused by dysregulation of NIX and BNIP3 turnover.

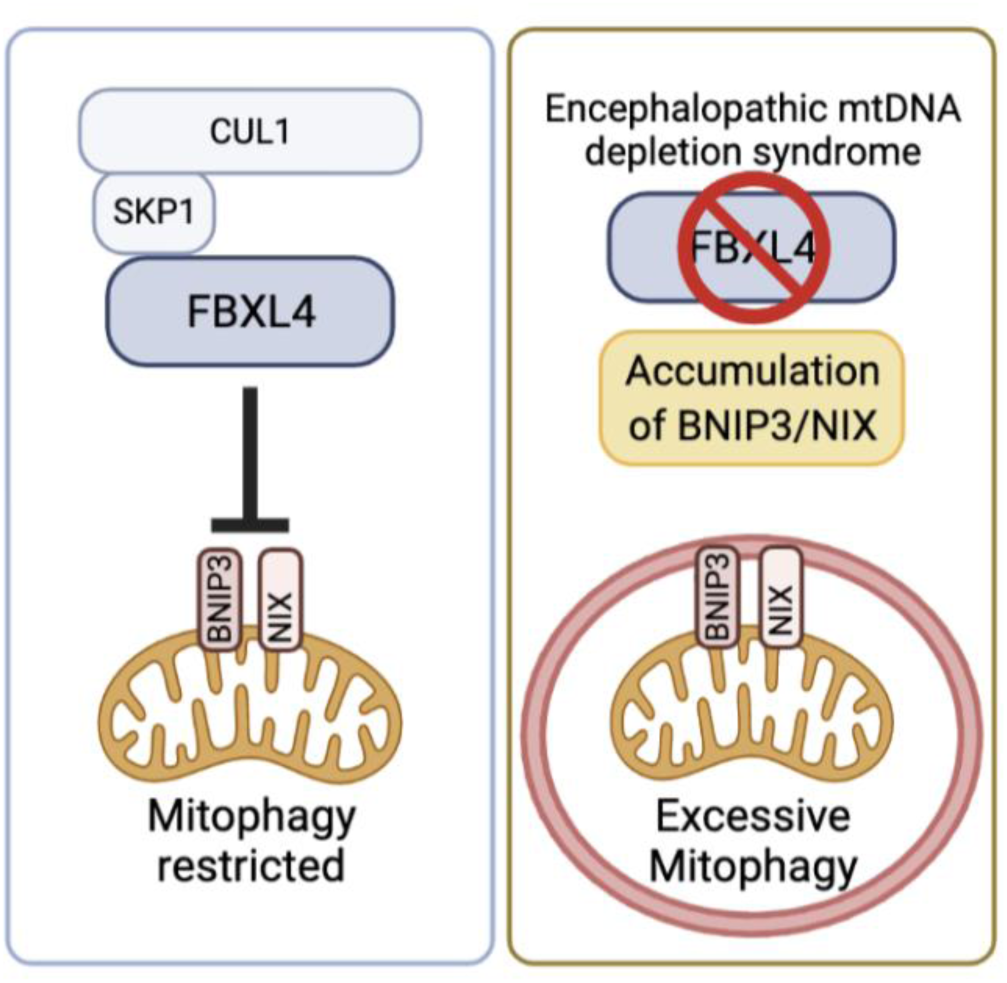

## Introduction

Mitophagy, otherwise known as mitochondrial autophagy, is the selective degradation of surplus, aged, or damaged mitochondria via autophagy. In this process, mitochondria are engulfed in a double membrane vesicle, the autophagosome, which ultimately fuses with the lysosome for degradation (Onishi *et al*, 2021; Pickles *et al*, 2018). Multiple distinct mitophagy pathways operate in response to a range of different mitochondrial stressors or physiological cues, including mitochondrial membrane depolarization (Jin *et al*, 2010; Narendra *et al*, 2008), hypoxia (Allen *et al*, 2013; Bellot *et al*, 2009; Sowter *et al*, 2001), or cellular differentiation (Esteban-Martinez *et al*, 2017; Sandoval *et al*, 2008; Schweers *et al*, 2007; Simpson *et al*, 2021). In addition, it is increasingly recognised that mitophagy occurs under basal conditions (i.e., in the absence of induced mitochondrial damage) (Lee *et al*, 2018; McWilliams *et al*, 2018). In contrast to stimulus-induced mitophagy, the mechanisms by which cells regulate mitophagy at steady state are not understood.

Mitophagy is initiated by specific signals on the mitochondrial outer membrane that are thought to serve as docking sites for the nascent autophagosome. Depolarization-induced mitophagy involves the Parkinson’s disease proteins, Pink1 and Parkin, which cooperate to induce the ubiquitylation of outer membrane proteins which indirectly induce autophagosome formation via autophagy adaptors (Lazarou *et al*, 2015). In contrast, hypoxia-induced or developmentally programmed mitophagy are triggered through the upregulation of mitophagy receptors NIX and BNIP3 that directly reside in the mitochondrial outer membrane (Bellot *et al*., 2009; Esteban-Martinez *et al*., 2017; Novak *et al*, 2010; Sandoval *et al*., 2008; Schweers *et al*., 2007; Simpson *et al*., 2021; Zhang *et al*, 2008). NIX and BNIP3 are approximately 50% homologous, and both contain an LC3-interaction motif (LIR), an atypical BH3 domain, and a C-terminal transmembrane domain (required for localization in the mitochondrial outer membrane). The LIRs of NIX and BNIP3 face the cytoplasm where they recruit the LC3 proteins on the autophagosome (Hanna *et al*, 2012; Novak *et al*., 2010). NIX and BNIP3 are barely detectable at mitochondria in unstressed conditions but are upregulated through HIF1α-mediated transcription in response hypoxia and iron chelation to mediate mitophagy (Allen *et al*., 2013; Sowter *et al*., 2001; Zhao *et al*, 2020).

Cullin-RING ligases (CRLs) comprise the largest family of multi-subunit E3 ligases (Harper & Schulman, 2021; Lydeard *et al*, 2013). Each CRL complex contains one of 8 different Cullin subunits, which act as assembly scaffolds, binding at their C-termini to a RING finger protein (RBX1 or RBX2), which is required for binding to the E2 ubiquitin conjugating enzyme. To recognize specific substrates, each CRL complex binds to adaptor proteins which recruit variable substrate recognition proteins at their N-termini. The SCF (SKP1-CUL1-F-box protein) sub-family of CRLs (also known as CRL1 complexes) consist of the CUL1 backbone, the RBX1 RING subunit, the adaptor protein SKP1, and one of 69 different F-box proteins in humans as a substrate binding component (Duan & Pagano, 2021; Skaar *et al*, 2013), one of which is the mitochondria-localized F-box protein, FBXL4.

In humans, pathogenic, bi-allelic *FBXL4* variants result in encephalopathic mitochondrial DNA (mtDNA) depletion syndrome (MTDPS13) (Ballout *et al*, 2019; Bonnen *et al*, 2013; Gai *et al*, 2013), a multi-system disease that presents with congenital lactic acidosis, neurodevelopmental delays, poor growth, and encephalopathy (Bonnen *et al*., 2013; Gai *et al*., 2013). FBXL4-deficiency leads to severe oxidative phosphorylation deficiency correlating with a quantitative loss of mtDNA copy number (mtDNA depletion), hyper-fragmentation of the mitochondrial network and diminished steady-state levels of mitochondrial proteins (Alsina *et al*, 2020; Ballout *et al*., 2019; Bonnen *et al*., 2013; Gai *et al*., 2013; Sabouny *et al*, 2019). Despite the serious consequences of FBXL4 deficiency, no mitochondrial substrates for FBXL4 have yet been identified.

Here, we report a mechanism whereby SCF-FBXL4 constitutively targets the mitophagy receptors NIX and BNIP3 for degradation, restricting steady-state mitophagy. We found that MTDPS13-associated pathogenic variants of FBXL4 are unable to efficiently mediate NIX and BNIP3 degradation. Our results suggest that the increased basal mitophagy and associated molecular phenotypes in FBXL4-associated mtDNA depletion syndrome are caused by NIX and BNIP3 hyperaccumulation.

## Results

HIF1α is the master regulator of hypoxia- and iron chelation-induced mitophagy via transcriptional upregulation of NIX and BNIP3 mitophagy receptors (Allen *et al*., 2013; Sowter *et al*., 2001; Zhao *et al*., 2020). This pathway is antagonized by the activity of the CRL2-VHL ubiquitin ligase, which mediates the polyubiquitylation and proteolytic degradation of HIF1α (Ivan *et al*, 2001; Jaakkola *et al*, 2001; Maxwell *et al*, 1999), thus suppressing mitophagy by preventing both HIF1α stabilization and the consequent upregulation of BNIP3 and NIX. We investigated whether, in addition to CRL2-VHL, other CRLs play a role in mitophagy regulation, possibly through targeting BNIP3 and NIX for degradation directly. To do this, we inhibited the entire CRL family using MLN4924, an inhibitor of Cullin Neddylation (Soucy *et al*, 2009), and removed the contribution of HIF1α using echinomycin, a HIF1α inhibitor (Kong *et al*, 2005). Mitophagy was assessed using the pH-sensitive mito-Keima (mt-Keima) reporter and confocal microscopy to detect mito-lysosomes (Sun *et al*, 2017). To avoid contributions from Parkin-mitophagy, U2OS or HeLa cells were used for mitophagy assays, with low or no Parkin expression, respectively (Munson *et al*, 2022; Tang *et al*, 2017).

Cells treated with either the iron chelator deferiprone (DFP) or MLN4924, which are both HIF1α stabilizers (Allen *et al*., 2013), displayed robust mitophagy (Figure 1A and 1B), correlating with the upregulation of NIX and BNIP3 (Figure 1C and EV1A-C, E). As expected, DFP-induced mitophagy was eliminated by echinomycin treatment (Zhao *et al*., 2020); however, in contrast, we found that MLN4924-induced mitophagy was only partially eliminated by echinomycin (Figure 1A and 1B), demonstrating that one or more CRL(s) suppresses mitophagy via a HIF1α-independent mechanism. Similarly, whilst echinomycin or depletion of HIF1α by siRNA prevented the DFP-induced increase in NIX and BNIP3 protein levels, it did not prevent the increase of NIX and BNIP3 protein levels in response to MLN4924 (Figure 1C and EV1C), demonstrating that one or more CRL(s) contributes to the turnover of NIX and BNIP3 protein levels. Lastly, using BNIP3/NIX double knockout (BNIP3/NIX DKO) cells, we established that mitophagy detected after MLN4924 treatment occurs through NIX and/or BNIP3 (Figure 1C, 1G, EV1B and Table EV1).

**Figure 1.**
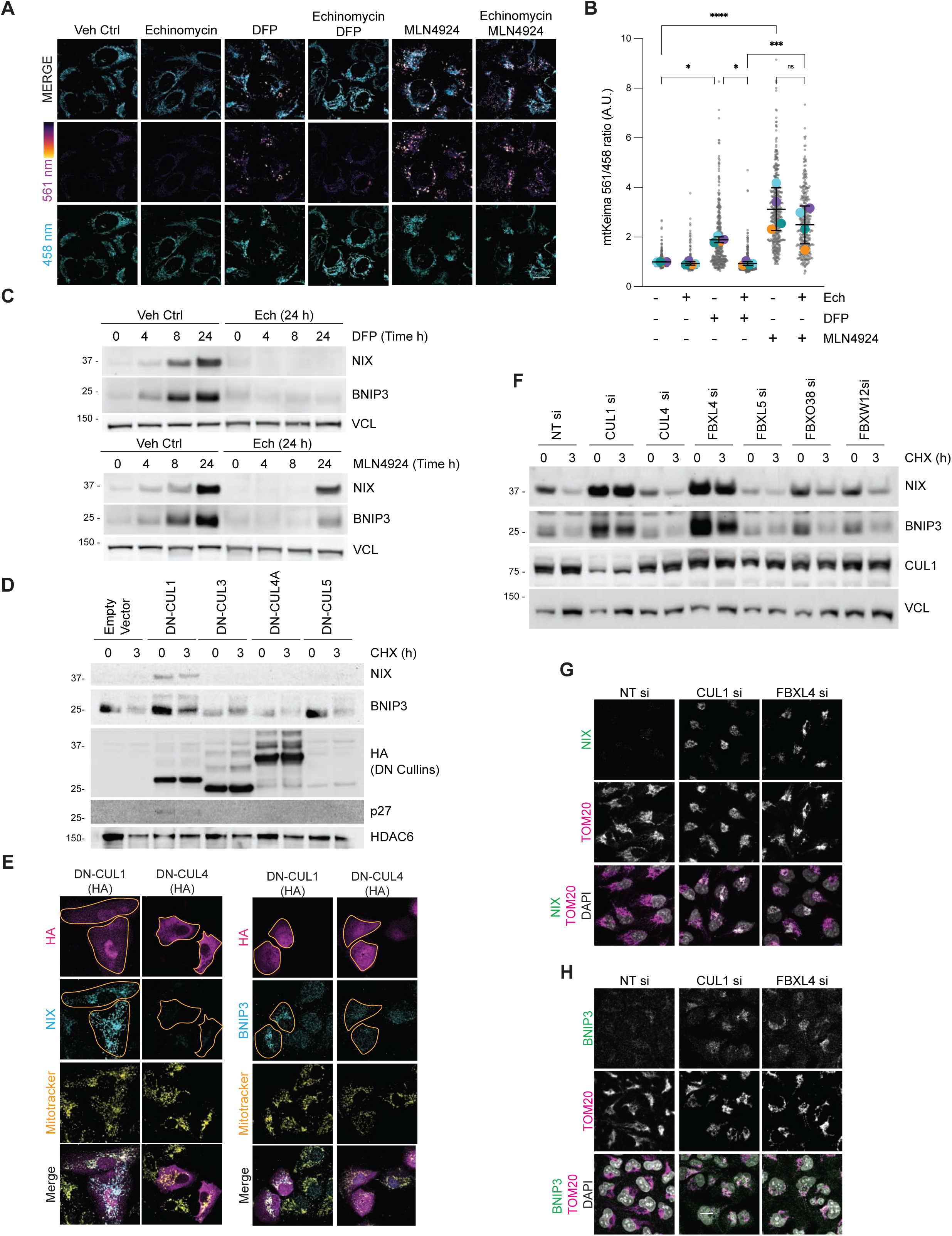
Identification of SCF-FBXL4 as a negative regulator NIX and BNIP3 stability. A) *Cullin-RING ligases suppress mitophagy in a HIF1α-independent manner.* Inhibition of HIF1α with echinomycin prevents DFP-induced mitophagy but does not prevent MLN4924-induced mitophagy. U2OS cells stably expressing mt-Keima were treated for 24 h, as indicated (DFP 1 mM; MLN4924 0.5 μM; Echinomycin 10 nM) and analyzed by live-cell confocal microscopy. The emission signal obtained after excitation with the 458 nm laser (neutral pH) or 561 nm laser (acidic pH) is shown in cyan or mpl inferno, respectively. B) *Quantification of A.* Mitophagy is represented as the ratio of mt-Keima 561 nm fluorescence intensity to mt-Keima 458 nm fluorescence intensity for individual cells normalised to the mean of the untreated condition. Translucent grey dots represent measurements from individual cells. Colored circles represent the mean ratio from independent experiments. The centre lines and bars represent the mean of the averaged independent replicates +/- standard deviation. N=3. C) *Cullin-RING ligase(s) negatively regulate NIX and BNIP3 protein levels in a HIF1α-independent manner*. Inhibition of HIF1α with echinomycin prevents the increase of NIX and BNIP3 in response to DFP, but not MLN4924. U2OS cells were treated with the indicated drugs for the specified times. Total-cell lysates were subject to immunoblotting as shown. D) *Expression of dominant-negative (DN) Cullin 1 results in an increase in the levels and half-life of NIX and BNIP3 protein.* HeLa-T-REx-Flp-in cells were transfected with FLAG-HA-tagged dominant-negative CUL1, CUL3, CUL4A and CUL5 or an empty vector, as indicated. 24 h post-transfection, cells were treated with cycloheximide for 3 h. Cell lysates were immunoblotted with the indicated antibodies. E) *Expression of dominant-negative (DN) Cullin 1 results in the accumulation of NIX and BNIP3 at mitochondria.* U2OS cells were transfected with FLAG-HA-tagged DN-CUL1 or FLAG-HA-tagged DN-CUL4 and immunostained for both HA and either NIX or BNIP3. An orange line marks the edge of the individual cells expressing the dominant-negative cullin protein. Scale bars = 10 μm. F) *Screening F-box proteins for changes in NIX and BNIP3 protein stability.* NIX and BNIP3 are stabilized by depletion of CUL1 and FBXL4 (but not other F-box proteins). U2OS cells were transfected with the indicated siRNAs. Total-cell lysates were subject to immunoblotting as shown. G) *Depletion of FBXL4 and CUL1 results in NIX accumulation at mitochondria.* U2OS cells were transfected with non-targeting siRNA, CUL1 siRNA or FBXL4 siRNA. Cells were fixed and stained with the indicated antibodies. H) *Depletion of FBXL4 and CUL1 results in BNIP3 accumulation at mitochondria.* U2OS cells were transfected with non-targeting siRNA, CUL1 siRNA or FBXL4 siRNA. Cells were fixed and stained with the indicated antibodies. P values were calculated based on the mean values from independent experiments using one-way ANOVA (\**P*<0.05, \*\**P*<0.005, \*\*\**P*<0.001, \*\*\*\**P*<0.0001). Scale bars = 20 μm.

Collectively, these results suggest that a CRL-based mechanism basally restricts mitophagy in cells, in a HIF1α-independent manner, possibly through post-translational regulation of BNIP3/NIX.

Next, to narrow down which cullin-RING ligase family is involved in turnover of NIX and BNIP3, we examined the effect of disrupting individual cullin proteins using dominant-negative (DN) versions. Expression of these proteins interferes with the function of the respective endogenous cullin, resulting in the accumulation of their specific substrates (Emanuele *et al*, 2011; Simoneschi *et al*, 2021). We transfected dominant-negative (DN) versions of CUL1, CUL3, CUL4A and CUL5 into cells, finding that the expression of DN-CUL1, but not other DN-cullin proteins, increased the steady-state levels and extended the half-lives of NIX and BNIP3 (Figure 1D). Furthermore, cells expressing DN-CUL1 displayed an accumulation of NIX and BNIP3 at the mitochondria when compared with either the surrounding untransfected cells or with cells expressing DN-CUL4 (Figure 1E). These findings indicate that NIX and BNIP3 mitophagy receptors are subject to SCF-ubiquitin ligase-mediated turnover.

CUL1 forms the backbone of 69 distinct SCF complexes, each containing a different F-box protein (Skaar *et al*., 2013). To identify the specific F-box protein(s) targeting NIX and/or BNIP3 to the SCF complex, we screened a partial siRNA library targeting F-box proteins for increased levels of NIX and BNIP3 (Figure 1F, shows 4 of 11 tested). Of the F-box proteins assessed, the depletion of FBXL4 resulted in the greatest upregulation of both NIX and BNIP3 protein levels. To test whether the half-life of NIX or BNIP3 is extended after depletion of FBXL4, we performed a cycloheximide-chase assay and found that silencing of FBXL4 or CUL1 promoted the stabilization of both NIX and BNIP3, whereas silencing of CUL4 did not (Figure EV2A). Similarly, the depletion of FBXL4 or CUL1, but not CUL4, resulted in the upregulation of NIX and BNIP3 at mitochondria (Figure 1G-H). In all, this data indicates that SCF-FBXL4 mediates the turnover of NIX and BNIP3 mitophagy receptors under steady-state conditions.

### FBXL4 localizes to the outer mitochondrial membrane and controls the ubiquitylation and turnover of NIX and BNIP3

To confirm that FBXL4 mediates the turnover of NIX and BNIP3, we generated FBXL4-deficient U2OS cell lines using CRISPR/Cas9-mediated gene disruption. The resulting FBXL4-deficient clones contained a frameshift mutation leading to an early termination codon at position Arg209 (for clone FBXL4-2G10) and a 5 amino acid deletion between Glu367-Glu372 (for clone FBXL4-1D4) (Table EV1). Corresponding with increased stability of NIX and BNIP3 as assessed by cycloheximide chase assay (Figure 2A), we found that both FBXL4-deficient cell lines displayed significantly higher levels of NIX and BNIP3 at mitochondria (Figure 2B and 2C). Rescue experiments demonstrated that inducible expression of FBXL4 tagged with HA at its C-terminus (FBXL4^HA-C^) in both FBXL4-deficient cell lines (FBXL4-2G10 and FBXL4-1D4) was able to restore the elevated NIX and BNIP3 protein levels back to parental levels, further demonstrating that FBXL4 mediates the turnover of NIX and BNIP3 (Figure EV2B and 3C). This downregulation of NIX and BNIP3 by FBXL4 required FBXL4’s mitochondria-localisation sequence (MTS, amino acids 1-29; see Figure EV4B for the localisation of FBXL4^Δaa1-29^) and a functional F-box domain (required to bind to SKP1 and CUL1 in the SCF core complex, see Figure 4C), indicating that FBXL4 activity depends on its mitochondrial localisation and its interaction with SKP1 and CUL1 (Figure 2D).

**Figure 2.**
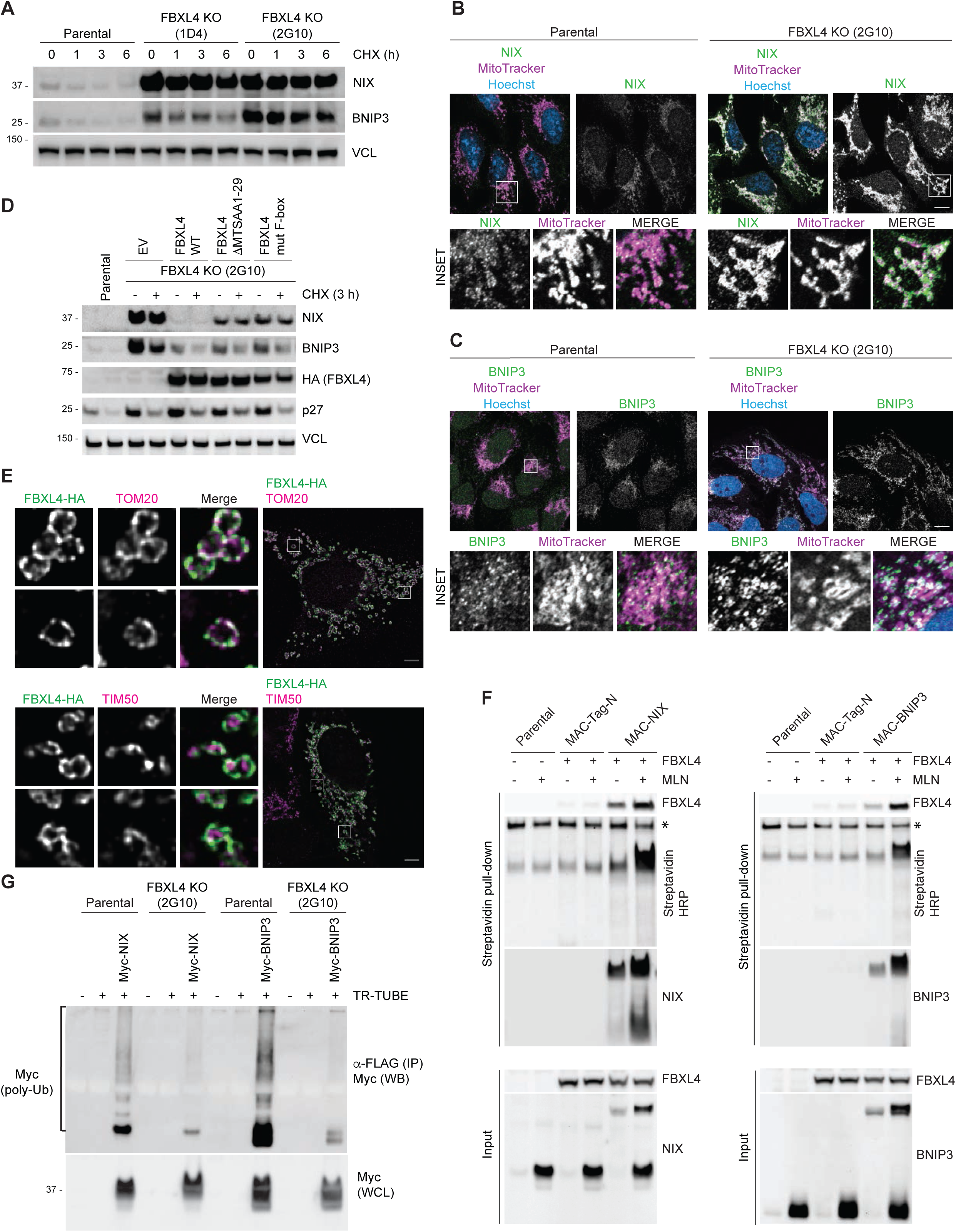
FBXL4 localizes to the mitochondrial outer membrane and controls the turnover and ubiquitylation of NIX and BNIP3. A) *NIX and BNIP3 are upregulated and stabilized in CRISPR-Cas9 generated FBXL4-deficient cells.* CRISPR-mediated genome editing was used to modify the *FBXL4* locus in U2OS cells. Clonal cell lines lacking FBXL4 were treated with cycloheximide for the indicated times prior to immunoblotting. B) *NIX accumulates at mitochondria in FBXL4-deficient cells.* FBXL4-deficient cells (clone 2G10) were fixed and stained with MitoTracker Red and with antibodies to NIX (green). Scale bar = 10 μm. C) *BNIP3 accumulates at mitochondria in FBXL4-deficient cells.* FBXL4-deficient cells (clone 2G10) were fixed and stained with MitoTracker Red and with antibodies to BNIP3 (green). Scale bar = 10 μm. D) *Re-expression of FBXL4 into FBXL4-deficient CRISPR lines destabilizes NIX and BNIP3 and this depends on FBXL4’s mitochondrial targeting sequence and F-box domain.* U2OS FBXL4 KO (2G10) cells were rescued with wild-type FBXL4-HA or variants lacking either the mitochondrial targeting sequence (FBXL4-ΔMTS) or the F-box domain (FBXL4-F-box mut) variants. Cells were treated with cycloheximide for 3 h prior to harvesting. E) *FBXL4 localizes to the mitochondrial outer membrane.* Cells transfected with FBXL4-HA-C were treated with DFP for 24 h. Cells were stained with an anti-HA-tag antibody (to recognize FBXL4) and either TOM20 (an outer mitochondrial membrane protein) or TIM50 (an inner mitochondrial membrane protein). Scale bar = 5 μm. F) *FBXL4 is a proximity interactor of NIX and BNIP3.* Cells expressing inducible BirA-BNIP3, BirA-NIX and BirA control were transduced with a lentiviral vector expressing FBXL4, as indicated. Cells were treated with doxycycline for 48 h (to induce BirA-bait protein expression), biotin for 24 h (for the biotinylation reaction), and, where indicated, MLN4924 for 24 h (to stabilise NIX and BNIP3). Streptavidin-coupled beads were used to capture the biotinylated proteins. FBXL4 was specifically detected in the eluate from BirA-BNIP3 and BirA-NIX, but not BirA-alone. *Non-specific band. G) *NIX and BNIP3 polyubiquitylation depends on FBXL4.* U2OS or U2OS-FBXL4 KO (2G10) cells were co-transfected with TR-TUBE and either myc-BNIP3 or myc-NIX, as indicated. Cell lysates obtained 48 h post-transfection were immunoprecipitated with anti-FLAG antibody, and the immunoprecipitates were analyzed by immunoblotting. The line on the left marks a ladder of bands corresponding to polyubiquitylated myc-BNIP3 or myc-NIX.

**Figure 3.**
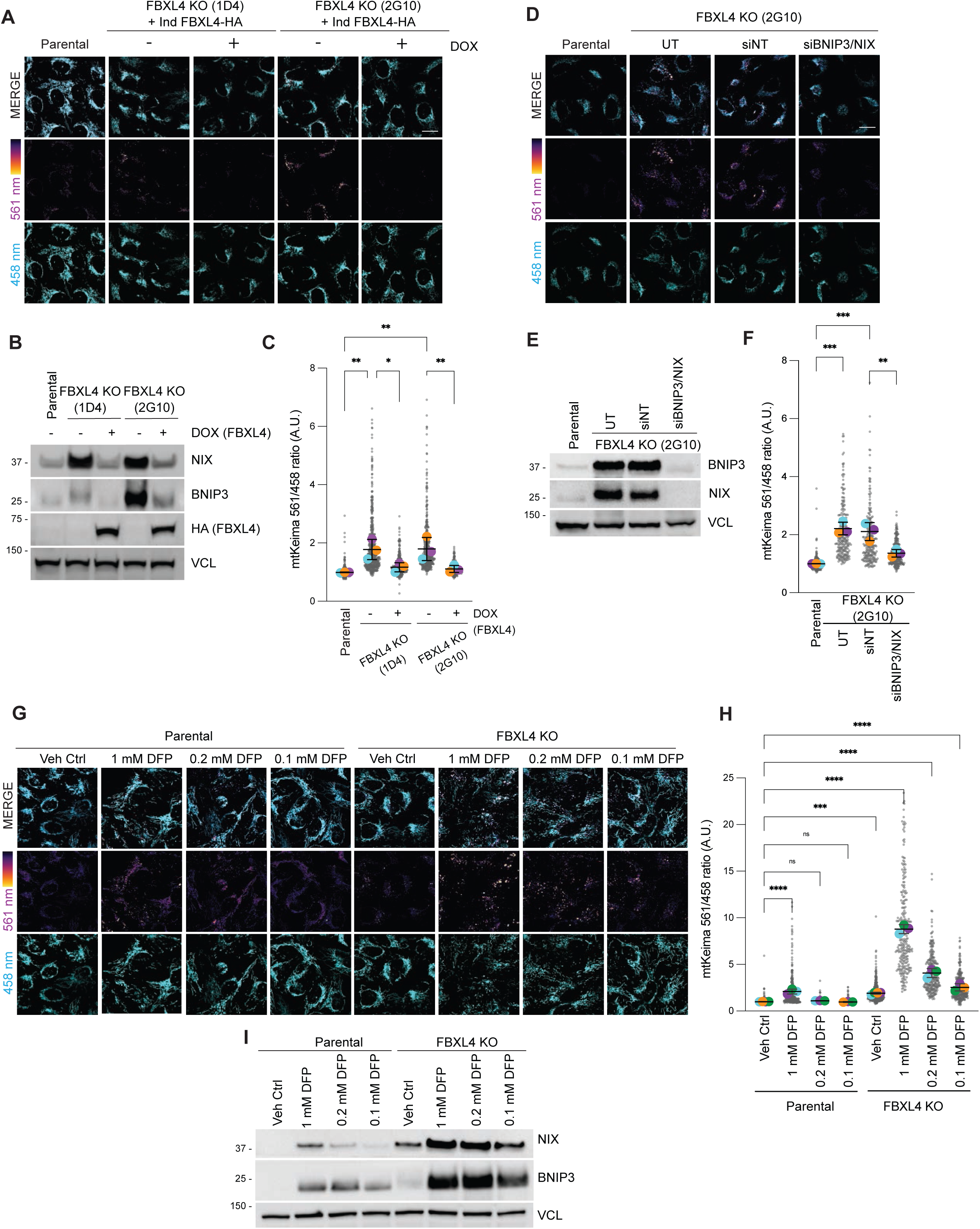
FBXL4-deficiency promotes mitophagy through BNIP3/NIX stabilization. A) *Mitophagy increases upon FBXL4 disruption and is rescued by the induction of HA-FBXL4.* U2OS mt-Keima FBXL4 KO clones (2G10 and 1D4) expressing doxycycline-inducible wild-type FBXL4-HA were treated with doxycycline for 72 h. The emission signal obtained after excitation with the 458 nm laser (neutral pH) or 561 nm laser (acidic pH) is shown in cyan or mpl inferno, respectively. B) *Quantification of A.* Mitophagy is represented as the ratio of mt-Keima 561 nm fluorescence intensity divided by mt-Keima 458 nm fluorescence intensity for individual cells normalised to the parental condition. Translucent grey dots represent measurements from individual cells. Colored circles represent the mean ratio from independent experiments. The centre lines and bars represent the mean of the independent replicate means +/- standard deviation. N=3. C) Corresponding cells from (3A) were harvested for immunoblotting to analyze the extent of NIX and BNIP3 reduction by induction of FBXL4-HA. D) *BNIP3/NIX depletion by siRNA reduces mitophagy after FBXL4 disruption.* U2OS mt-Keima cells and U2OS mt-Keima FBXL4 KO 2G10 cells were transfected with siRNAs targeting both NIX and BNIP3 (siBNIP3/NIX) or non-targeting siRNA (siNT). Live-cell confocal microscopy was performed to visualise mitophagy. UT=untransfected. E) *Quantification of D.* N=3. F) Corresponding cells from (3D) were harvested for immunoblotting to analyze the extent of NIX and BNIP3 reduction by siRNA. G) *FBXL4-deficient cells are ultra-sensitive to DFP-induced mitophagy.* U2OS mt-Keima cells and U2OS mt-Keima FBXL4 KO 2G10 were treated with DFP at specified concentrations for 24 h and analyzed by live-cell confocal microscopy. H) Quantification of G. N=3. I) Cells from (3G) were lyzed and immunoblotting was performed. P values were calculated from the mean values from independent experiments using one-way ANOVA (\**P*<0.05, \*\**P*<0.005, \*\*\**P*<0.001, \*\*\*\**P*<0.0001). Scale bars = 20 μm.

**Figure 4.**
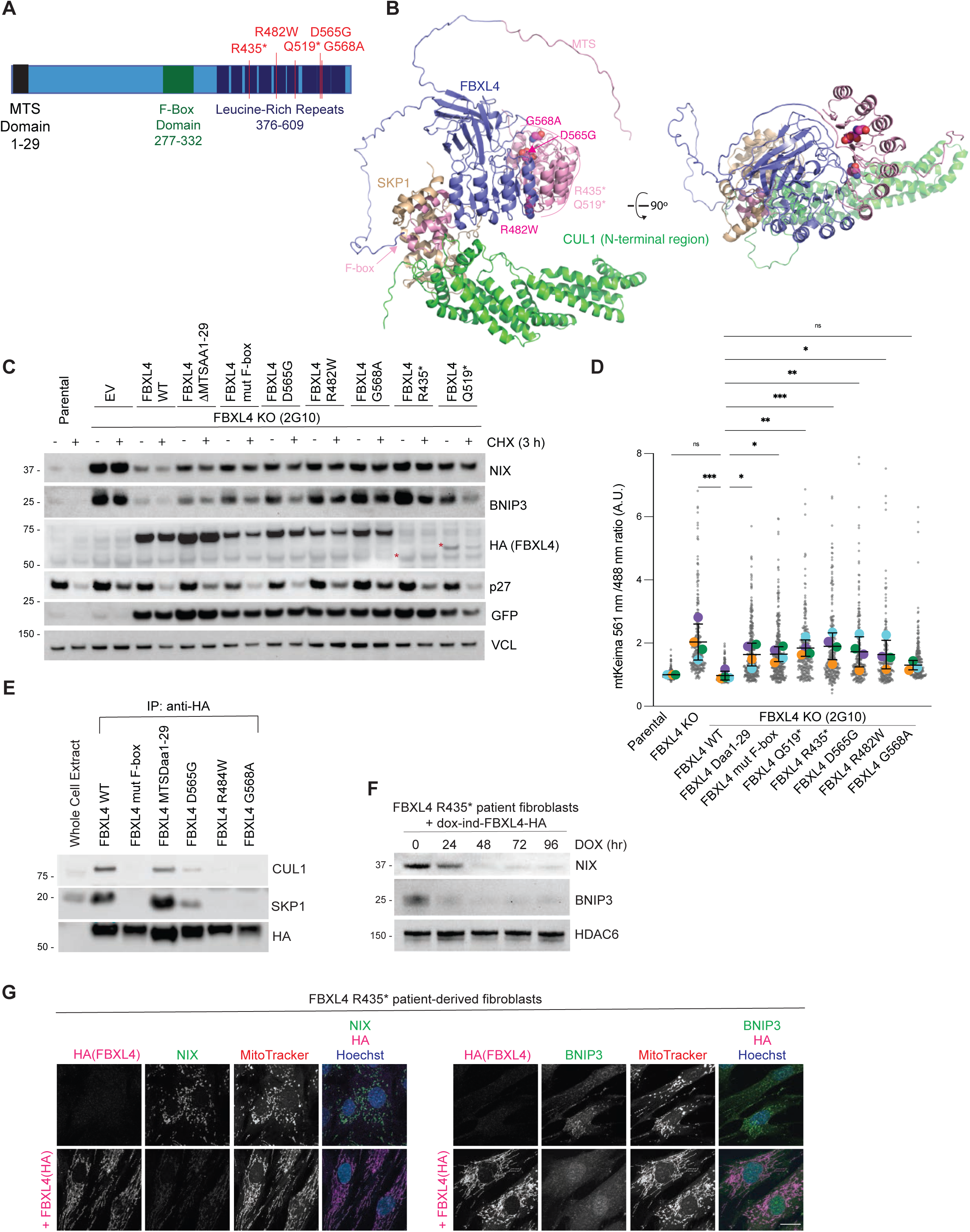
MTDPS13 patient-derived FBXL4 variants do not efficiently assemble into an SCF complex have impaired abilities to mediate NIX and BNIP3 turnover. A) *Schematic representation of domain structure of FBXL4.* Pathological variants tested herein are shown in red. B) *Alphafold2 structural modelling of FBXL4 and its complex formation with SCF components SKP1 and CUL1.* Pathogenic variants of FBXL4 indicated in magenta spheres. The pale pink section of the LRRs represents the region deleted by the truncation deletions (Arg435). C) *FBXL4 patient variants are less efficient than wild-type FBXL4 in mediating NIX and BNIP3 downregulation and destabilization.* U2OS FBXL4 KO (2G10) cells were rescued with constructs expressing wild-type FBXL4-HA, FBXL4(F-box mut), FBXL4(ΔMTSaa1-29) or specified patient variants. Cells were treated with cycloheximide for 3 h prior to harvesting. Samples were lyzed, and immunoblotting was performed. GFP serves as a marker of transduction efficiency/transgene expression. EV = empty vector. D) *FBXL4 patient variants are less efficient than FBXL4-wild-type at suppressing mitophagy.* U2OS mt-Keima cells, U2OS mt-Keima FBXL4 KO 2G10 cells and 2G10 cells rescued with the specified FBXL4 constructs were analyzed by confocal microscopy to quantify mitophagy. Mitophagy is represented as the ratio of mt-Keima 561 nm fluorescence intensity divided by mt-Keima 458 nm fluorescence intensity for individual cells normalised to untreated U2OS cells. Translucent grey dots represent measurements from individual cells. Colored circles represent the mean ratio from independent experiments. The centre lines and bars represent the mean of the independent replicates +/- standard deviation. P values were calculated based on the mean values using a one-way ANOVA (*P<0.05, **P<0.005, ***P<0.001, ****P<0.0001). N=4. E) *FBXL4-Arg482Trp, FBXL4-Asp565Gly, FBXL4-Gly568Ala patient variants are less efficient than FBXL4-wild-type at assembling into a complex with SKP1 and CUL1.* FBXL4-KO cells expressing wild-type FBXL4-HA or FBXL4 variants were harvested and lyzed. Whole-cell extracts were subjected to immunoprecipitation (IP) with anti-HA agarose beads and immunoblotting, as indicated. F) *Reconstitution of FBXL4-HA into FBXL4-deficient patient fibroblast cells causes down-regulation of NIX and BNIP3.* FBXL4-deficient patient fibroblasts (derived from patients harboring homozygous non-sense mutation in *FBXL4* at pArg435*) were transduced with doxycycline-inducible FBXL4-HA constructs. Cells were harvested at the indicated times following doxycycline induction. Immunoblotting was performed as indicated. G) *Reconstitution of FBXL4-HA into FBXL4-deficient patient fibroblast cells causes down-regulation of NIX and BNIP3.* FBXL4-deficient patient fibroblasts were transduced with FBXL4-HA. Cells were stained with MitoTracker, fixed and co-immunostained with antibodies to HA (to detect FBXL4) and either NIX or BNIP3. Scale bar = 20 μm

NIX and BNIP3 have C-terminal transmembrane domains and localize to the mitochondrial outer membrane (Figure EV3I). To determine if FBXL4 also localizes to the mitochondrial outer membrane allowing it to promote the turnover of NIX and BNIP3, we analyzed the localization of FBXL4^HA-C^. Our initial attempts to distinguish between outer membrane and inner membrane localization were not definitive due to the small diameter of filamentous mitochondria in U2OS cells. We therefore treated cells with DFP, which promotes the formation of swollen and donut-shaped mitochondria, to allow us to distinguish between mitochondrial substructures more easily. We found that, in most cells, FBXL4^HA-C^ colocalized with TOM20, an outer mitochondrial membrane protein, and on the outside of TIM50, an inner membrane protein (Figure 2E and EV2C). This led us to conclude that FBXL4 predominantly localises to the mitochondrial outer membrane.

To further demonstrate spatial proximity between FBXL4 and both NIX and BNIP3, we used proximity-dependent biotin identification (BioID). BioID uses a biotin ligase BirA, fused to a bait (in this case either NIX or BNIP3), to biotinylate prey proteins within a ∼10-nm labelling radius. BioID was chosen for its utility in the detection of interactions under denaturing conditions (allowing for the solubilisation of mitochondrial membrane proteins) as well as in the detection of weak or transient interactions prior to cell lysis. Cells stably expressing inducible BirA-tagged BNIP3 or BirA-tagged NIX were incubated with biotin and biotinylated proteins were captured using Streptavidin-coupled beads. In addition, we treated cells with MLN4924 to prevent CRL-dependent degradation of NIX and BNIP3. Immunoblot analysis of the biotinylated proteins isolated in the Streptavidin-pulldown revealed both BirA-BNIP3 and BirA-NIX, but not BirA-alone control, associated with FBXL4^HA-C^ (Figure 2F), indicating that FBXL4 is co-located with NIX and BNIP3.

We next tested whether NIX and BNIP3 are ubiquitylated in an FBXL4-dependent manner. To this end, we co-transfected U2OS cells with myc-tagged BNIP3 or myc-tagged NIX together with FLAG-tagged TR-TUBE (Yoshida *et al*, 2015) and investigated whether they coimmunoprecipitated. TR-TUBE is a tandem ubiquitin-binding entity that directly binds polyubiquitin chains and protects them from proteasome-mediated degradation and thus can detect ubiquitylated substrates. Immunoprecipitation of TR-TUBE showed a co-precipitating smear of high-molecular weight species of myc-BNIP3 and myc-NIX reflecting their polyubiquitylation. The ubiquitylated species induced by TR-TUBE and detected using the anti-myc antibody were dramatically reduced in the FBXL4 knockout cell line, indicating that ubiquitylation of both NIX and BNIP3 relies on the presence of FBXL4 (Figure 2G).

### FBXL4-deficiency promotes mitophagy through BNIP3/NIX stabilization

To test the hypothesis that the elevated levels of NIX and BNIP3 in FBXL4-deficient cells are sufficient to induce mitophagy in basal conditions, we used the mt-Keima mitophagy assay described in Figure 1. FBXL4-deficient U2OS cell lines exhibited increased mitophagy compared with parental cell lines and this was rescued by re-introducing FBXL4 (Figure 3A-C). Importantly, the elevated mitophagy in FBXL4-deficient cells was reduced when NIX and BNIP3 were depleted by siRNA, supporting that FBXL4 restricts BNIP3- and NIX-dependent mitophagy (Figure 3D-F). Next, we examined if FBXL4 suppresses DFP-induced mitophagy. We found that FBXL4-deficient cells treated with DFP (at increasing concentrations) exhibited substantially enhanced mitophagy (and NIX and BNIP3 levels) compared to parental cells treated with DFP, suggesting that the elevated levels of mitophagy receptors sensitises cells to DFP (Figure 3G-I).

To directly determine the effect of NIX and BNIP3 stabilization on mitophagy, we sought to generate hyper-stable versions of NIX and BNIP3 mutants. Unlike F-box proteins that recognise short degrons on their substrates (e.g. βTRCP), crystal structures of F-box proteins of the LRR family (e.g. FBXL3) suggest that their interaction with substrates can occur over large surfaces (Skaar *et al*., 2013; Xing *et al*, 2013), precluding the mapping of short degron sequences that disrupt binding to FBXL4. Therefore, to identify the respective regions in NIX and BNIP3 required for their destabilization, we generated a series of deletion constructs for inducible expression in HeLa T-rex Flp-in cells and identified stable mutants based on their higher levels and longer half-lives in the presence of cycloheximide (Figure EV3A-EV3D). We identified several adjacent highly conserved C-terminal regions in NIX and BNIP3 that, when deleted, resulted in their stabilisation compared with wild-type NIX and BNIP3. Specifically, the region of aa 151-184 in NIX and aa 161-225 in BNIP3 contributed to their destabilization (Figure EV3C and EV3D, respectively).

To test the hypothesis that stabilization of NIX or BNIP3 mimics FBXL4 deficiency, we next induced the expression of the hyper-stable mutants of NIX or BNIP3 in mt-Keima-expressing HeLa cells. In the absence of any external mitophagic triggers, we found that the inducible expression of hyper-stable NIXΔ150-171 or NIXΔ170-184 resulted in approximately 2-fold increase in the mean mitophagy ratio compared with wild-type NIX (Figure EV3E-F). Likewise, the inducible expression of BNIP3 stable mutants also triggered mitophagy when compared to wild-type BNIP3 (Figure EV2G-H). Notably, all stable deletion mutants localized normally to mitochondria (Figure EV3I).

Taken together, these results suggest that FBXL4 suppresses mitophagy by restraining the levels of NIX and BNIP3 mitophagy receptors.

### MTDPS13 patient-derived FBXL4 variants have impaired abilities to mediate NIX and BNIP3 turnover and restrict mitophagy

We tested whether the pathogenic *FBXL4* variants responsible for MTDPS13 (OMIM # 615471) interfere with FBXL4 function. FBXL4 possesses a typical F-box domain that associates directly with SKP1, a C-terminal LRR (leucine-rich repeat) domain consisting of twelve repeats, and a unique N-terminal β-sheet domain with a nine-stranded discoidin-like fold (Figure 4A, 4B and EV4A). The N-terminal domain of FBXL4 is not found in other FBXL family members and is predicted to form an intimate intramolecular interaction with the C-terminal LRR domain (Figure 4B). Most of the pathogenic mis-sense variations in *FBXL4* are in its C-terminal LRR domain (the putative substrate binding region). To test whether pathological FBXL4 variants are as efficient as wild-type FBXL4 at mediating NIX and BNIP3 turnover, we performed rescue experiments in FBXL4-deficient U2OS cells complemented with either wild-type FBXL4 or MTDPS13-associated FBXL4 variants (Arg482Trp, Asp565Gly, Gly568Ala, Gln519*, and Arg435*, based on RefSeq <u>NM_001278716.2</u>). Whereas wild-type FBXL4 was able to reduce the levels and half-lives of NIX and BNIP3 to basal levels, the disease-associated FBXL4 variants displayed an impaired ability to promote NIX and BNIP3 turnover (Figure 4C). The Gln519-term and ArgR435-term truncation variants were expressed at significantly lower levels than wild-type FBXL4, however FBXL4 missense variants (Arg482Trp, Asp565Gly, Gly568Ala) were expressed at levels similar to wild-type FBXL4, suggesting that their inability to degrade NIX and BNIP3 was not related to their expression levels. Correlating with their inability to mediated NIX and BNIP3 turnover, FBXL4 pathogenic variants were also less effective at suppressing mitophagy when reconstituted into FBXL4-deficient cells (Figure 4D and EV4B). Notably, despite their reduced function, the FBXL4 variants localized, like wild-type FBXL4, to mitochondria (Figure EV4C).

To explore how the disease-associated FBXL4 variants affect NIX and BNIP3 degradation, we next examined the ability of FBXL4 variants to bind to SKP1 and CUL1, core members of the SCF complex. Although the disease-associated variants are in the LRR region outside the F-box domain, we found that none of the variants was as effective as wild-type FBXL4 at binding to SKP1 and CUL1 (Figure 4E). Mutations that affect substrate binding could impair SCF-FBXL4 complex assembly, as is the case for FBXL3 which only assembles an SCF complex in the presence of substrate (Yumimoto *et al*, 2013). Alternatively, the mutations could broadly affect protein folding and in that way impede SCF assembly.

Finally, to determine if NIX and BNIP3 accumulate in patients with MTDPS13, we examined NIX and BNIP3 levels in a fibroblast cell line derived from a patient homozygous for the p.Arg435* *FBXL4* variant (Alsina *et al*., 2020; Bonnen *et al*., 2013). NIX and BNIP3 levels were readily detectable in this cell line but were downregulated when wild-type FBXL4-HA was reintroduced (Figure 4F-G).

Taken together, our results suggest that MTDPS13-associated pathogenic FBXL4 variants have impaired abilities to mediate the degradation of NIX and BNIP3 mitophagy receptors, resulting in their accumulation and increased mitophagy.

## Discussion

The cellular triggers promoting basal mitophagy are poorly understood. Here, we demonstrate that FBXL4 restricts the abundance of NIX and BNIP3 mitophagy receptors to suppress mitophagy in basal conditions. Thus, NIX and BNIP3 are negatively regulated by two distinct CRLs: 1) the CRL2-VHL complex which mediates the turnover of HIF1α and thereby inhibits NIX and BNIP3 transcription, and 2) the SCF-FBXL4 complex at the mitochondrial outer membrane. Our data reveal that mitophagy is actively suppressed by the continuous degradation of NIX and BNIP3, previously thought to be upregulated solely at the level of transcription. The multiple mechanisms converging to regulate the abundance of NIX and BNIP3 have presumably evolved to ensure tight regulation of mitophagy levels for the cell to respond precisely and rapidly to changes in metabolic signals.

Dysregulation of FBXL4 function results in mtDNA depletion syndrome 13. Despite the serious consequences of *FBXL4* mutations (Alsina *et al*., 2020; Bonnen *et al*., 2013; Gai *et al*., 2013), the molecular functions of the FBXL4 protein have remained elusive (i.e., no mitochondrial substrates for FBXL4 have been previously identified). Our data uncover a mechanistic link between FBXL4 and mitophagy in MTDPS13, demonstrating that MTDPS13-derived FBXL4 variants are defective in mediating the turnover NIX and BNIP3 mitophagy receptors.

How FBXL4 activity is regulated remains to be elucidated. It is an interesting prospect that FBXL4 localization or activity could be inhibited on specific mitochondria selected for mitophagy to allow NIX and BNIP3 accumulation. Unlike the transcriptional regulation of NIX and BNIP3 via HIF1α, such local regulation of FBXL4 (and thus NIX and BNIP3) would represent a mechanism to control turnover of selected mitochondria, rather than global pools of mitochondria that are removed by mitophagy for metabolic re-programming in response to hypoxia (Zhang *et al*., 2008). Similarly, post-translational modifications on NIX and BNIP3 that stabilize them through disrupting their recognition by SCF^FBXL4^ may occur on specific mitochondria, allowing selective targeting.

Notably, NIX and BNIP3 accumulate indiscriminately (i.e., non-selectively) on the outer membrane of all mitochondria upon loss of FBXL4. However, only a proportion of tagged mitochondria undergo mitophagy despite the stabilisation of NIX and BNIP3, implying that additional signalling or stochastic events contribute to mitophagy induction. How mitophagy receptor stabilization cooperates with other signalling mechanisms and the fission-fusion machinery to facilitate mitophagy is unclear. Multiple mechanisms have been reported to facilitate mitophagy induction via mitophagy receptors, e.g., phosphorylation (Chen *et al*, 2014; Liu *et al*, 2012; Rogov *et al*, 2017; Wu *et al*, 2014) and dimerization (Marinkovic *et al*, 2021).

Future investigations will focus on understanding the precise mechanism by which FBXL4 recognises NIX and BNIP3. Although we did not yet identify the interface through which FBXL4 engages NIX and BNIP3, we were able to identify regions within NIX and BNIP3 that when deleted, resulted in stabilisation of these proteins and consequently increased mitophagy in the absence of overt mitochondrial stress. Whether these broad regions in the C-termini of NIX and BNIP3 represent sites of potential sites of post-translational modifications that support ligase recognition or access to the sites of ubiquitylation remains to be determined.

## Materials and methods

### Antibodies

Mouse monoclonal anti-TOM20 (clone 29; 612278) and mouse monoclonal anti-p27 (clone 57/Kip1/p27; 610242) were obtained from BD Biosciences. Mouse monoclonal anti-TIM50 (clone C-9; sc-393678), mouse monoclonal anti-BNIP3 (clone ANa40; sc-56167), mouse monoclonal anti-NIX (clone H-8; sc-166332), mouse monoclonal anti-vinculin (VCL, clone G-11; sc-55465), mouse monoclonal anti-γ−Tubulin (clone C-11; sc-17787), rabbit polyclonal anti-HDAC6 (sc-11420), and mouse monoclonal anti-GFP (B-2, sc-9996) were obtained from Santa Cruz Biotechnology. Mouse monoclonal anti-HA (clone 16B12; 901513) was obtained from BioLegend. Rabbit monoclonal anti-BNIP3 (clone EPR4034; ab109362) was obtained from Abcam. Mouse monoclonal anti-Myc (clone 9B11; 2276S), mouse monoclonal anti-HA Alexa Fluor^TM^ 488 conjugate (clone 6E2; 2350S), rabbit monoclonal anti-NIX (clone D4R4B;12396), rabbit monoclonal anti-HA (clone C29F4; 3724S), rabbit monoclonal anti-LC3B (clone D11; 3868S) and rabbit monoclonal anti-HIF1α (clone D1S7W; 36169S) were obtained from Cell Signaling Technology. Rabbit polyclonal anti-CUL1 (718700) was obtained from Thermo Fisher Scientific. Mouse monoclonal anti-FLAG (clone M2; F3165) and rabbit polyclonal anti-FLAG (SAB4301135) were obtained from Sigma-Aldrich. Rabbit anti-SKP1 was generated in the Pagano laboratory (Pagan *et al*, 2015). Secondary donkey anti-mouse IgG Alexa Fluor^TM^ 488 (A21202), donkey anti-mouse IgG Alexa Fluor ^TM^ 555 (A31570), donkey anti-mouse IgG Alexa Fluor ^TM^ 594 (A21203), donkey anti-mouse IgG Alexa Fluor ^TM^ 647 (A31571), donkey anti-rabbit IgG Alexa Fluor^TM^ 488 (A21026), donkey anti-rabbit IgG Alexa Fluor^TM^ 555 (A31572), donkey anti-rabbit IgG Alexa Fluor^TM^ 647 (A31573) were obtained from Thermo Fisher Scientific. Goat anti-rabbit IgG Atto 647N (40839) was purchased from Sigma-Aldrich.

### DNA constructs

pCHAC-mt-mKeima was a gift from R. Youle (RRID: Addgene 72342) (Lazarou *et al*., 2015). pLIX_402 was a gift from David Root (Addgene plasmid # 41394). MAC (BirA-Ha-Strep-tag II)-N was a gift from Markku Varjosalo (Addgene plasmid # 108078). FLAG-tagged TR-TUBE has been previously published (Yoshida *et al*., 2015). pcDNA5/FRT/TO/FLAG-S-tag has been previously published (Pagan *et al*., 2015). Dominant-negative Cullin constructs, including pcDNA3-Flag-HA-DN-CULLIN1(1-252), pcDNA3-Flag-HA-DN-CULLIN3(1-240), pcDNA3-Flag-HA-DN-CULLIN4(1-237), and pcDNA3-Flag-HA-DN-CULLIN5(1-228) were generated by site-directed mutagenesis. The pcDNA3.1(+)-N-Myc-BNIP3, pcDNA3.1(+)-N-Myc-NIX, pDONR-N-FLAG-BNIP3, pDONR-N-FLAG-BNIP3Δ141-160, pDONR-N-FLAG-BNIP3Δ161-192, pDONR-N-FLAG-BNIP3Δ193-225, pDONR-N-FLAG-BNIP3Δ181-203, pDONR-N-FLAG-NIX, pDONR-N-FLAG-NIXΔ120-150, pDONR-N-FLAG-NIXΔ151-170, pDONR-N-FLAG-NIXΔ171-184, pcDNA3.1(+)-C-HA-FBXL4, pcDNA3.1-C-eGFP-FBXL4, pDONR-C-HA-FBXL4 were generated by Genscript®. pLV-FBXL4-C-HA:IRES:EGFP, pLV-FBXL4-C-HA-F-BOX mut(LP283AA;LP297AA):IRES:EGFP, pLV-FBXL4-C-HA-ΔMTS(Δ1-29):IRES:EGFP, pLV-FBXL4-C-HA(Asp565Gly):IRES:EGFP, pLV-FBXL4-C-HA(Arg482Trp):IRES:EGFP, pLV-FBXL4-C-HA(Gly568Ala):IRES:EGFP, pLV-FBXL4-C-HA(Gly519 term):IRES:EGFP, pLV-FBXL4-C-HA(Arg435 Term):IRES:EGFP were generated by VectorBuilder. The Gateway cloning system (Thermo Fisher Scientific) was used to generate pcDNA5/FRT/TO/FLAG and pLIX-402 based constructs.

### CRISPR/Cas9-mediated genome editing

The pSpCas9 BB-2A-Puro (PX459) plasmid backbone was used to create the following guide RNA (gRNA) plasmids (created by Genscript®): BNIP3 CRISPR gRNA plasmid (gRNA targeting sequence: TCTTGTGGTGTCTGCGAGCG), NIX CRISPR gRNA plasmid (gRNA targeting sequence: TAGCTCTCAGGTGTGTCGGG); and FBXL4 CRISPR gRNA plasmids (gRNA targeting sequence: CAATTCAAGGCGTACTAATT; gRNA targeting sequence 2: CCCCACAAATCTTATACGAC).

To generate CRISPR/Cas9 knockout (KO) cell lines, cells were transiently transfected with the CRISPR gRNA plasmids targeting the gene(s) of interest. Twenty-four h post-transfection, cells were selected with puromycin (Sigma) for 72 h. They were then diluted as one cell per well into 96-well plates until single colonies formed. Successful editing was screened for by immunoblot analysis and/or indirect immunofluorescence microscopy. Sanger sequencing was used to confirm the presence of frameshift indels in the potential KO clones first identified by immunoblotting or immunofluorescence screening. For this, genomic DNA was isolated using the salting out method (Miller *et al*, 1988). In brief, cells were lysed in lysis buffer (50mM Tris HCL, SDS 1%) and genomic DNA was precipitated following the adding of 5M NaCl, Proteinase K, and absolute ethanol. Then, PCR was performed to amplify the targeted regions. The PCR product was subcloned into pCR^TM^-BluntII-TOPO^®^ vector (Zero Blunt^®^ TOPO^®^ PCR cloning Kit, Invitrogen^TM^) and sequenced with M13 forward primer to characterize the indels (described in Appendix Table 1).

To validate BNIP3 knockout clones, a set of primers including BNIP3 forward (FWD) (5’- GAGGAAGAGTTTGGCTCTGGCAGG-3’) and BNIP3 reverse (RVS) (5’- CGGTGTATCCCTGATGGCAG-3’) was used. To validate NIX KO clones, a set of primers including NIX FWD (5’-AGTGCAGAACATTTTGGGAGT-3’) and NIX RVS (5’- AAATCACCCGTCTTCTGCGT-3’) was used. To check FBXL4 KO clones, two sets of primers including FBXL4 FWD (Guide1- 5’TTTTAGCCTAACCATTCATATTTCA-3’ or Guide2- 5’- CCTTAAGGGACCAGTAGATCTCA-3’) and FBXL4 RVS (Guide1- 5’CTGCCAGCATTTTGGCTTAC-3’ or Guide2- 5’-CAATGCTCAATTACCGATGC-3’) were used.

### Cell culture and chemicals

Cell lines were grown at 37°C in a humidified incubator containing 5% CO_2_. HeLa cells (ATCC CCL-2), U2OS (ATCC HTB-96) and HEK293T (ATCC CRL-3216) cells were maintained in Dulbecco’s modified Eagle’s medium/nutrient mixture F-12 GlutaMAX^TM^ (DMEM/F-12; Thermo Fisher Scientific) supplemented with 10% fetal bovine serum. Fibroblast cells derived from a patient with homozygous p.Arg435* *FBXL4* have been previously published (Bonnen *et al*., 2013) (Alsina *et al*., 2020) and were cultured in DMEM/F-12 GlutaMAX^TM^ with 20% FBS and 5 mg/mL penicillin and streptomycin (Thermo Fisher Scientific). Where indicated, cells were treated with cycloheximide (CHX; 100 µg/mL; 66-81-9), Deferiprone (DFP; 1 mM; 379409), DMOG (0.5 mM; D3695) and Echinomycin (10 nM; SML0477), which were purchased from Sigma. MLN4924 (0.5 µM; 85923S) was obtained from Cell Signaling Technology. MG132 (10 µM; 474787) was purchased from Merck.

### Cell line generation

FLAG-S-tagged BNIP3(WT), BNIP3(Δ141-160), BNIP3(Δ161-192), BNIP3(Δ193-225), and BNIP3(Δ181-203), NIX(WT), NIX(Δ120-150), NIX(Δ151-170), NIX(Δ171-184), MAC-N, MAC-BNIP3 and MAC-NIX were cloned into pcDNA5/FRT/TO (Thermo Fisher). Constructs were co-transfected with pOG44 into HeLa-T-rex Flp-in cells to generate inducible cell lines using Flippase (Flp) recombination target (FRT)/Flp-mediated recombination technology in HeLa-T-rex Flp-in cells, as previously described (Pagan *et al*., 2015). Twenty-four h post-transfection, cells were selected with Hygromycin B (400 μg/ml) for approximately ten days. HeLa-T-rex Flp-in cell-lines were subsequently maintained in Hygromycin B (200 μg/ml). To induce expression, cells were treated with 0.5 μg/mL doxycycline (Sigma; 10592-13-9).

To generate stably transfected cell lines, retrovirus (pCHAC-mt-mKeima) and lentiviruses (for pLix402 and pLV constructs) were packaged in HEK293T cells. HeLa or U2OS cells were transduced with virus for 48 h with 10 μg/ml polybrene (Sigma), then optimized for protein expression via fluorescence sorting or puromycin selection.

### Transfection

Plasmid transfections were performed using Lipofectamine 2000 (Thermo Fisher Scientific) and siRNA transfections were performed using Lipofectamine RNAiMAX (Thermo Fisher Scientific), as per manufacturer’s instructions. ON-TARGETplus Non-targeting Control Pool (Dharmacon; D-001810-01) was used as the siRNA control. ON-TARGETplus siCUL1 pool (L-004086-00), siCUL2 pool (L-007277-00), siCUL3 pool (L-010224-00), siCUL4 pool (L-012610-00), siCUL5 pool (L-019553-00), siFBXL4 pool (L-013564-00), siHIF1α pool (L-004018-00), siFBXL5 pool (L-012424-00), siFBXO38 pool (L-018163-00), siFBXW12 pool (L-032001-00), siBNIP3 pool (M-004636-01-0005) and siNIX pool (M-11815-01-0005) were purchased from Dharmacon^TM^ (Horizon Discovery).

### Immunoblotting

Immunoblotting was performed as previously described (Pagan *et al*., 2015). In brief, cells were harvested and subsequently lyzed in SDS lysis buffer (50 mM Tris and 2% SDS) at 97°C for 15 mins. Protein extracts were quantified using Direct Detect® Assay-free Cards (Merck; DDAC00010) or Pierce Bicinchoninic Bcid (BCA) assay (Thermo Fisher Scientific; 23250) and prepared for gel electrophoresis in Bolt™ LDS Sample Buffer (Invitrogen^TM^; B0008). Equal amounts of protein samples were resolved on SDS-PAGE (BOLT pre-cast 4-12% gradient gels, Invitrogen^TM^) and transferred onto methanol-activated Immobilon^®^-P PVDF Membrane (0.45 μm pore size) (Merck; IPVH00010) using BOLT gel transfer cassettes and BOLT transfer buffer (Invitrogen™; BT0006), according to the manufacturer’s instructions. The membranes were blocked in 5% skim milk for 1 h at room temperature and then incubated with indicated primary antibodies at 4°C overnight and secondary peroxidase-conjugated goat anti-rabbit or goat anti-mouse antibodies for 1 h at room temperature. The chemiluminescence signal was acquired using Pierce ECL Western blotting substrate (Thermo Fisher Scientific; 32106) or Pierce SuperSignal West Femto Substrate (Thermo Fisher Scientific; 34094) and ChemiDoc™ Imaging System (Bio-Rad).

### Co-immunoprecipitation assays

Cells were lyzed in a Tris-Triton lysis buffer (50 mM Tris-Cl pH 7.5, 150 mM NaCl, 10% glycerol, 1 mM EDTA, 1 mM EGTA, 5 mM MgCl_2,_ 1 mM β-glycerophosphate and 1% Triton) containing protease inhibitor cocktail (Rowe Scientific; CP2778) and PhosSTOP EASYpack Phosphatase Inhibitor Cocktail (Roche; 4906837001) on ice for 30 minutes. Cell lysates were collected by centrifugation at 14,000 rpm for 10 minutes at 4°C. To immunoprecipitate exogenously expressed FLAG-tagged or HA-tagged proteins, cell lysates were incubated in a rotating incubator for 1 h at 4°C with bead-conjugated FLAG (Sigma; A2220), bead-conjugated HA (Thermo Fisher Scientific; 88837), respectively. The immunoprecipitates were washed with Tris-Triton lysis buffer 5 times prior to elution with Bolt™ LDS Sample Buffer and Western blotting.

### BioID pulldown

Stable cells expressing doxycycline-inducible MAC(BirA-HA-Strep-tagII)-BNIP3, MAC-NIX, or MAC-N were generated and subsequently transduced with pLV-FBXL4-C-HA. Cells grown in 10 cm dishes were treated with 50 μM Biotin for 24 h. Cell pellets were lyzed in RIPA lysis buffer (50 mM Tris-HCl pH 7.5, 150 mM NaCl, 1% NP-40, 1mM EDTA, 1 mM EGTA, 0.1% SDS, protease inhibitors, and 0.5% Sodium deoxycholate) at 4°C for 1 h on a rotator. Lysates were sonicated (2 x 10 second bursts with 2 seconds rest in between) on ice at 50% amplitude. Lysates were then centrifuged for 30 min at 13,000 rpm at 4°C. Biotinylated proteins were captured using Pierce Streptavidin Magnetic Beads (Thermo Fisher Scientific, 88817) at 4°C on a rotator for 3 h. Magnetic beads collected on magnet for 1 minute between wash steps. The magnetic beads were washed with RIPA buffer (minus deoxycholate) 4 times prior to elution with 25 mM biotin at 95°C.

### TUBE Assay to detect polyubiquitylated proteins

Cells grown in 10 cm dishes were transiently transfected with 5 μg of FLAG-tagged TR-TUBE (Yoshida *et al*., 2015) and 5 μg of myc-tagged BNIP3 or myc-tagged NIX. For immunoaffinity purification of ubiquitylated proteins, cells were lyzed in Tris-Triton lysis buffer (50 mM Tris-Cl pH 7.5, 150 mM NaCl, 10% glycerol, 1 mM EDTA, 1 mM EGTA, 5 mM MgCl_2,_ 1 mM β-glycerophosphate and 1% Triton) and harvested 48 h post-transfection. Whole-cell lysates were incubated for 1 h with ANTI-FLAG^®^ M2 Affinity Gel (Sigma; A2220) followed by extensive washing. Bead-bound proteins were eluted using Bolt™ LDS Sample Buffer.

### Indirect Immunofluorescence

Adherent cells on coverslips were fixed in ice-cold methanol for 10 min at −20°C (for most antibodies) or fixed in 4% PFA for 1 h (for BNIP3-ANa40 antibody). Fixed cell monolayers were blocked with 2% BSA in PBS for 30 min to reduce non-specific binding. Cells were then sequentially labelled with diluted primary antibodies and corresponding secondary antibodies for 1 h at room temperature. Coverslips were mounted on glass microscope slides using Fluorescent Mounting Medium (Dako; S3023) or Prolong Diamond Antifade Mountant (Thermo Fisher Scientific; P36965). Images in Figure 2G and EV3I were acquired at room temperature using a DeltaVision Elite inverted microscope system (GE Healthcare) using a ×100/1.4NA Oil PSF Objective from Olympus. Optical sections were processed using the SoftWorx deconvolution algorithm. Images in Figure EV1E, EV2C-D were acquired using a Leica DMi8 SP8 Inverted confocal microscope equipped with 63x Plan Apochromatic objective. STED images in Figure EV2E were acquired using a Leica SP8 STED 3X Confocal Laser Scanning Microscope equipped with a 93x Plan Apochromatic objective and STED depletion lasers. The acquired images were then processed and exported using Huygens Deconvolution software. Images in Figure 1D, 2C and 2D and EV4A, were acquired using a Zeiss LSM900 Fast AiryScan2 Confocal microscope with a 63x C-Plan Apo NA 1.4 oil-immersion objective. Image deconvolution was performed using ZEN Blue 3D software (version 3.4).

### Protein structural prediction, modelling and visualisation

The structural predictions of human FBXL4 (Q9UKA2) was performed using the AlphaFold2 neural-network (Jumper *et al*, 2021) implemented within the freely accessible ColabFold pipeline (Mirdita *et al*, 2022). For each modelling experiment ColabFold was executed using default settings where multiple sequence alignments were generated with MMseqs2 (Mirdita *et al*, 2019) to produce five separate models per structure that were then subjected to energy minimisation with Amber (Eastman *et al*, 2017). In this instance, we verified that AlphaFold2 would produce a reliable predicted complex of the FBXL4 adaptor bound to SKP1 and the N-terminal region of CUL1. For producing images, structures were rendered with Pymol (Schrodinger, USA; https://pymol.org/2/).

### mt-Keima assay

The mt-Keima assay was performed as previously described (Sun *et al*., 2017). Dual-excitation (561/458 nm) images were acquired using a Leica DMi8 SP8 Inverted confocal microscope equipped with a 63x Plan Apochromatic objective and environmental chamber (set to 5% CO_2_ and 37°C). Quantitative analysis of mitophagy with mt-Keima was performed with Image J/Fiji software. Single cells were segregated from fields of view by generating regions of interest (ROI). The selected ROI was cropped and split into separate channels, prior to threshold processing. The fluorescence intensity of mt-Keima 561 nm (lysosomal signal) and mt-Keima 458 nm (mitochondrial signal) at the single-cell level were measured and the ratio 561 nm/458 nm was calculated. Three biological replicates were performed for each experiment, with >50 cells analyzed per condition for each repeat.

### Statistical analysis

Data were analyzed and using GraphPad Prism 9.0 software. The centre lines and error bars on graphs represent the mean of the averaged independent replicates +/- standard deviation. P values were calculated using one way ANOVA with post-hoc multiple comparisons, as described in the figure legends.

---

**Table S2.**
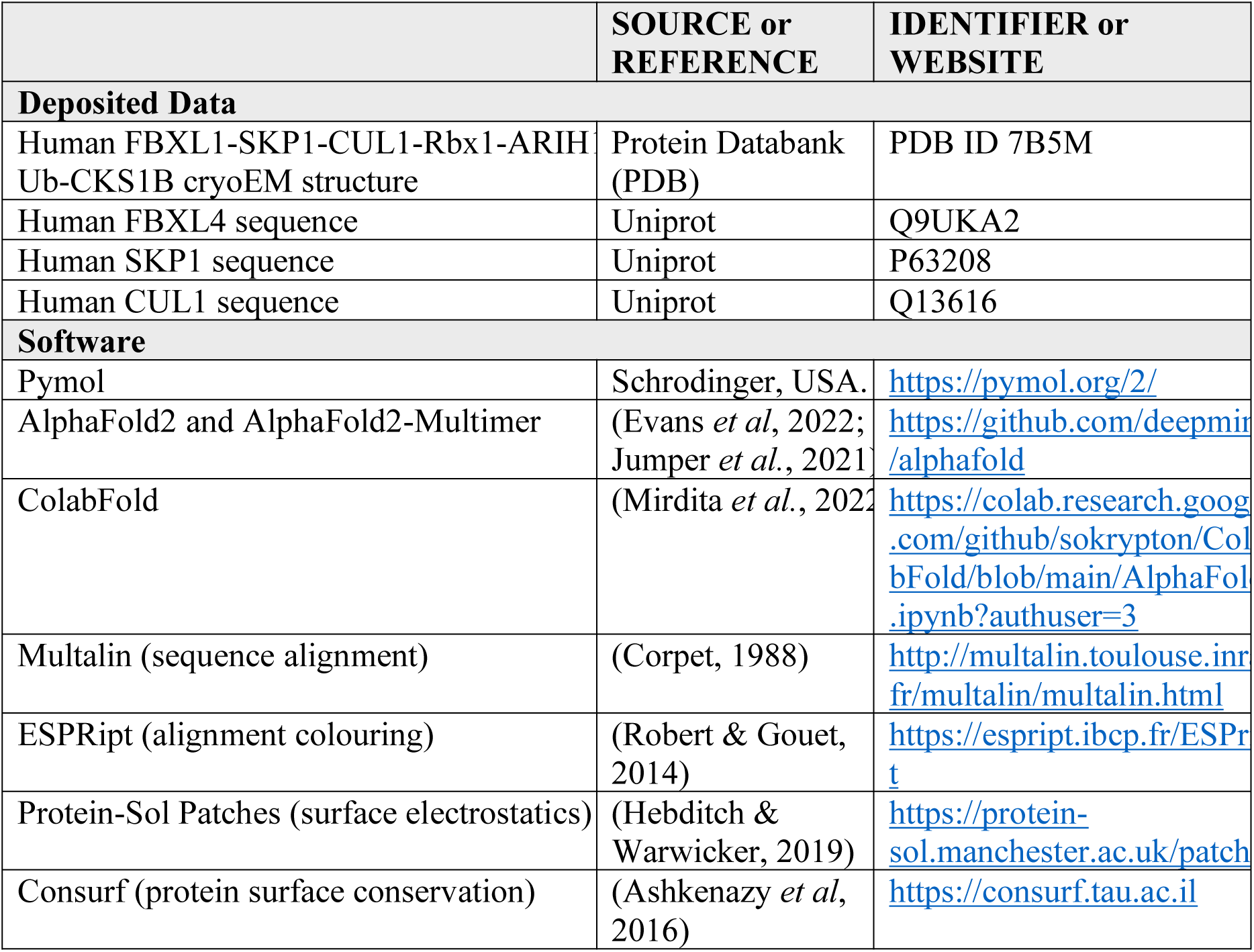

## Acknowledgements

We thank Rowan Tweedale, Stefan Thor and Uli Siebeck for critical reading and comments on the manuscript. We thank X.Qi for technical assistance. Imaging was performed at the Microscopy and Image Analysis Facility in the School of Biomedical Sciences, the University of Queensland. Lentiviruses were produced by the University of Queensland (UQ)-Viral Vector Core. This work was supported by an Australian National Health and Medical Research Council (APP1183915), Brain Foundation Research grant (2020), a Mito Foundation Incubator Grant (2022), and an Australian Research Council Future Fellowship (FT180100172) to J.K.P; National Health and Medical Research Council of Australia grants (APP1140064 and APP1150083 and fellowship APP1156489) to R.G.P.; an Australian Research Council Discovery Project (DP210102704) to M.J.K.J; an AIRC/Marie Curie, American Italian Cancer Foundation (AICF) and NIH/T32CA009161 grant to A.M. R.W.T. is funded by the Wellcome Centre for Mitochondrial Research (203105/Z/16/Z), the Mitochondrial Disease Patient Cohort (UK) (G0800674), the Medical Research Council International Centre for Genomic Medicine in Neuromuscular Disease (MR/S005021/1), the Medical Research Council (MR/W019027/1), the Lily Foundation, the Pathological Society, the UK NIHR Biomedical Research Centre for Ageing and Age-related disease award to the Newcastle upon Tyne Foundation Hospitals NHS Trust and the UK NHS Highly Specialised Service for Rare Mitochondrial Disorders of Adults and Children. B.M.C. is supported by an NHMRC Senior Research Fellowship (APP1136021) and an ARC Discovery Project Grant (APP1181135). SZ is supported by a Stafford Fox Foundation Fellowship. We would also like to acknowledge and thank Milot Mirdita, Sergey Ovchinnikov, Martin Steinegger and the ColabFold team for making their AlphaFold2 modelling pipeline available for public use. The authors declare that they have no conflict of interest.

## Contributions

G.N-D., K.S, Y.C. designed and performed most of the experiments. B.T., P.K., S.O., A.M., and M.J. assisted with experiments. Y.C., S.M., M.J, and J.P. supervised the students conducting the research. R.W.T. provided research materials and cell lines. M.L, S.Z. R.G.P., R.W.T., S.Z., B.C., and M.P. aided in analysis and interpretation of results. B.C. performed the Alphfold2 modelling. All authors discussed the results and commented on the manuscript. J.K.P. conceived and coordinated the study, oversaw the results, and wrote the manuscript.

---

**Figure EV1.**
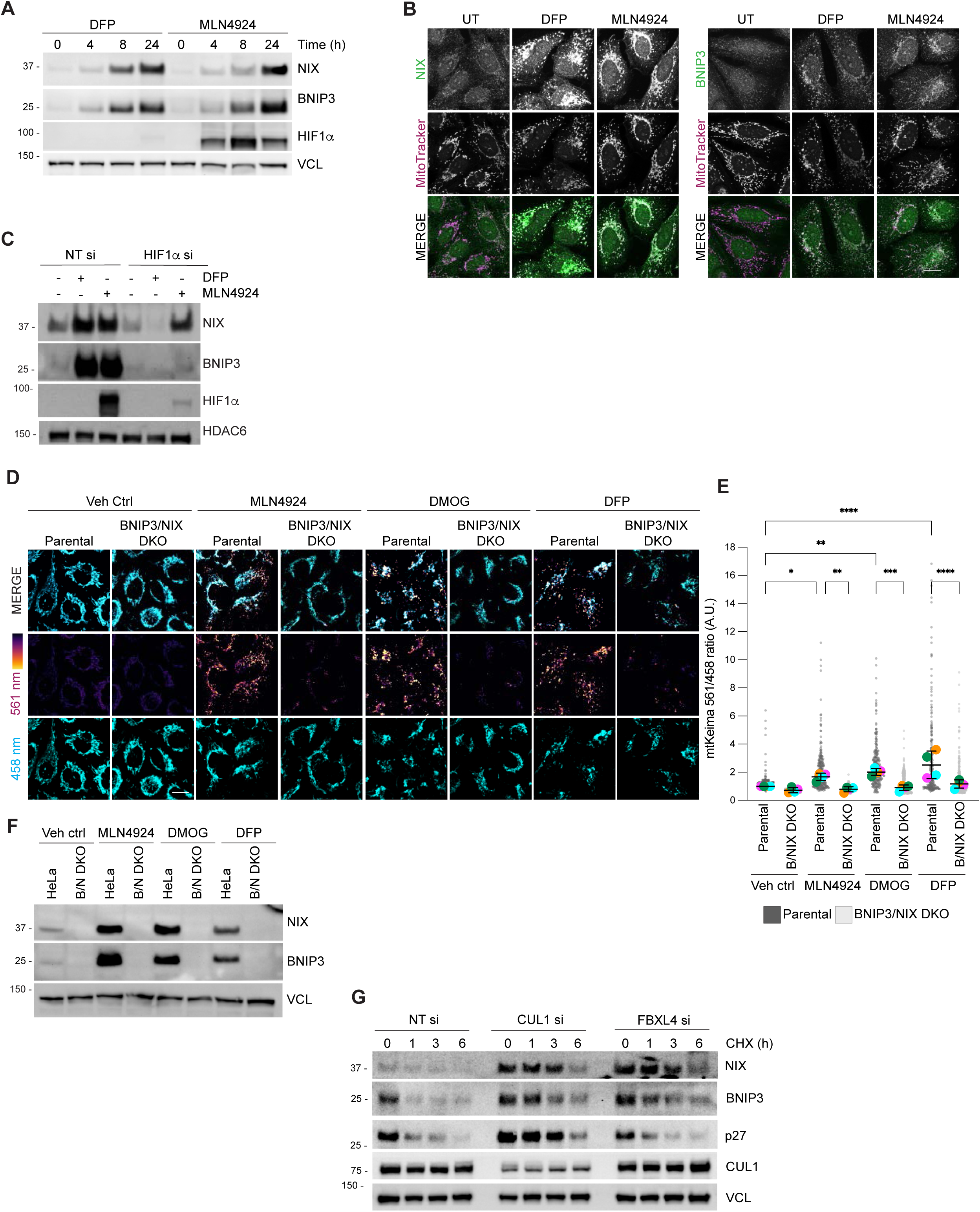
Identification of SCF-FBXL4 as a negative regulator NIX and BNIP3 stability. A) *NIX and BNIP3 protein levels increase in response to DFP or MLN4924 treatment.* U2OS cells were incubated with DFP (1 mM) or MLN4924 (0.5 μM) for the indicated times. Total cell lysates were subject to immunoblotting as shown. B) *NIX and BNIP3 protein levels increase at mitochondria in response to DFP or MLN4924 treatments.* U2OS cells were treated with DFP or MLN4924 for 24 h, fixed and stained with the indicated antibodies and MitoTracker Red. C) *Depletion of HIF1α with siRNA prevents the increase of NIX and BNIP3 in response to DFP, but not MLN4924.* U2OS cells were transfected with non-targeting siRNAs (NT si) or siRNAs targeting HIF1*α*. Cells were treated with DFP or MLN4924 for 24 h prior to harvesting for immunoblotting. D) *MLN4924-induced mitophagy requires NIX and BNIP3.* Parental HeLa or BNIP3/NIX double knockout (BNIP3/NIX DKO) HeLa cells expressing mt-Keima were treated for 24 h, as indicated. Cells were analyzed by live cell confocal microscopy, as described in A. E) *Quantification of D* was performed as described in B. Dark grey translucent dots represent measurements from individual parental HeLa cells. Light grey translucent dots represent measurements from individual BNIP3/NIX DKO cells. N=4. F) Analysis of NIX and BNIP3 protein levels in HeLa parental or BNIP3/NIX double knockout (BNIP3/NIX DKO) cells after MLN4924, DMOG, or DFP treatment. Total-cell lysates were subject to immunoblotting as shown. G) *NIX and BNIP3 are upregulated and stabilized by depletion of FBXL4 and CUL1.* U2OS cells were transfected with non-targeting siRNA, CUL1 siRNA or FBXL4 siRNA. Cells were treated with cycloheximide for the indicated time prior to immunoblotting with the specified antibodies.

**Figure EV2.**
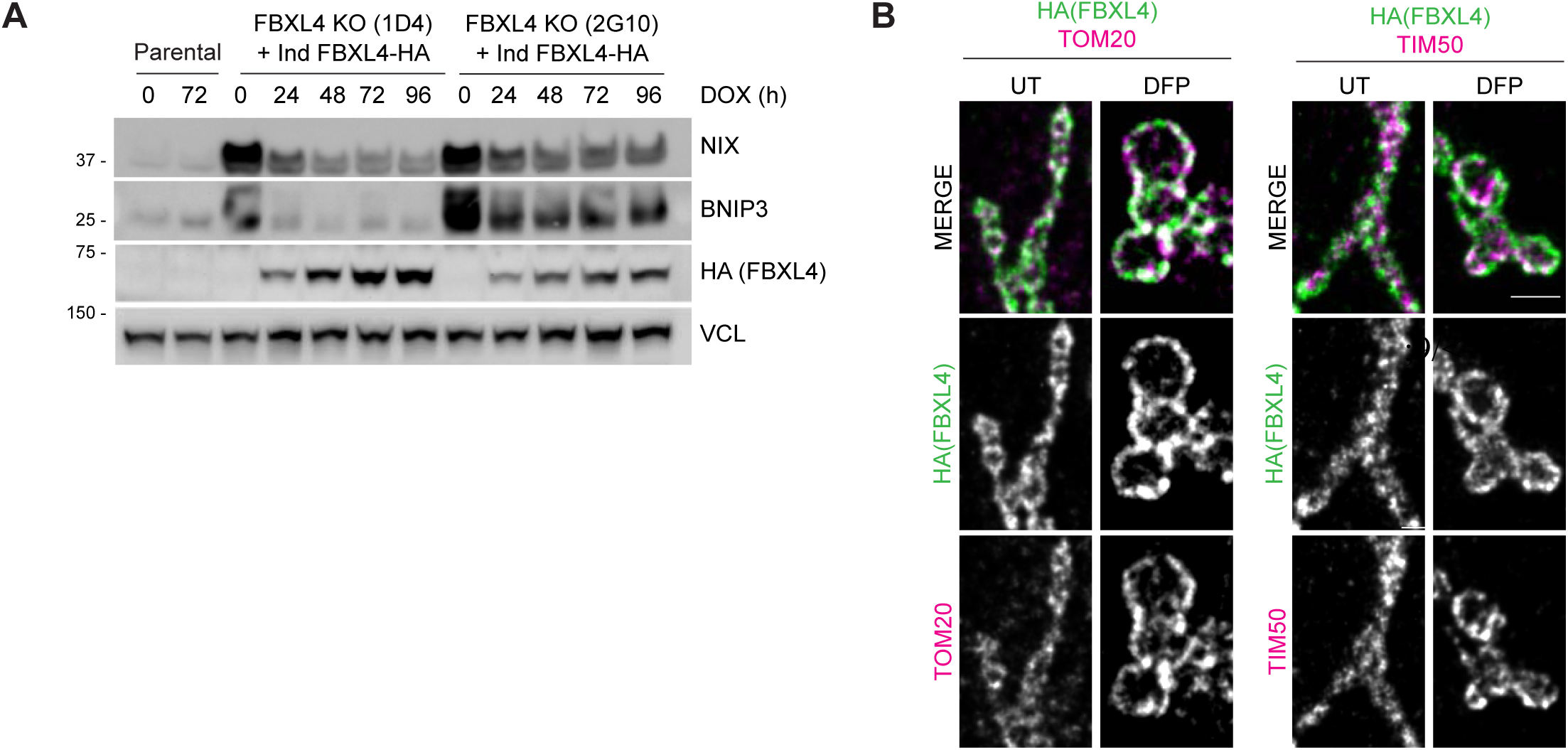
FBXL4 localizes to the mitochondrial outer membrane and controls the turnover and ubiquitylation of NIX and BNIP3. A) *Re-expression of FBXL4 into FBXL4-defective CRISPR lines rescues the levels of NIX and BNIP3 in multiple FBLX4 deficient clones.* FBXL4-deficient 2G10 and FBXL4-deficient 1D4 cell lines were stably transduced with a doxycycline-inducible FBXL4-HA construct. Cells were treated with doxycycline for the indicated times prior to immunoblotting with the specified antibodies. B) *FBXL4 localizes to the mitochondrial outer membrane.* U2OS cells transfected with FBXL4-HA-C were untreated (UT) or treated with DFP for 24 h. Two colour STED super-resolution microscopy of mitochondria was performed with anti-HA-tag antibody (to recognize FBXL4) and either TOM20 (an outer mitochondrial membrane protein) or TIM50 (an inner mitochondrial membrane protein). Scale bar = 1 μm.

**Figure EV3.**
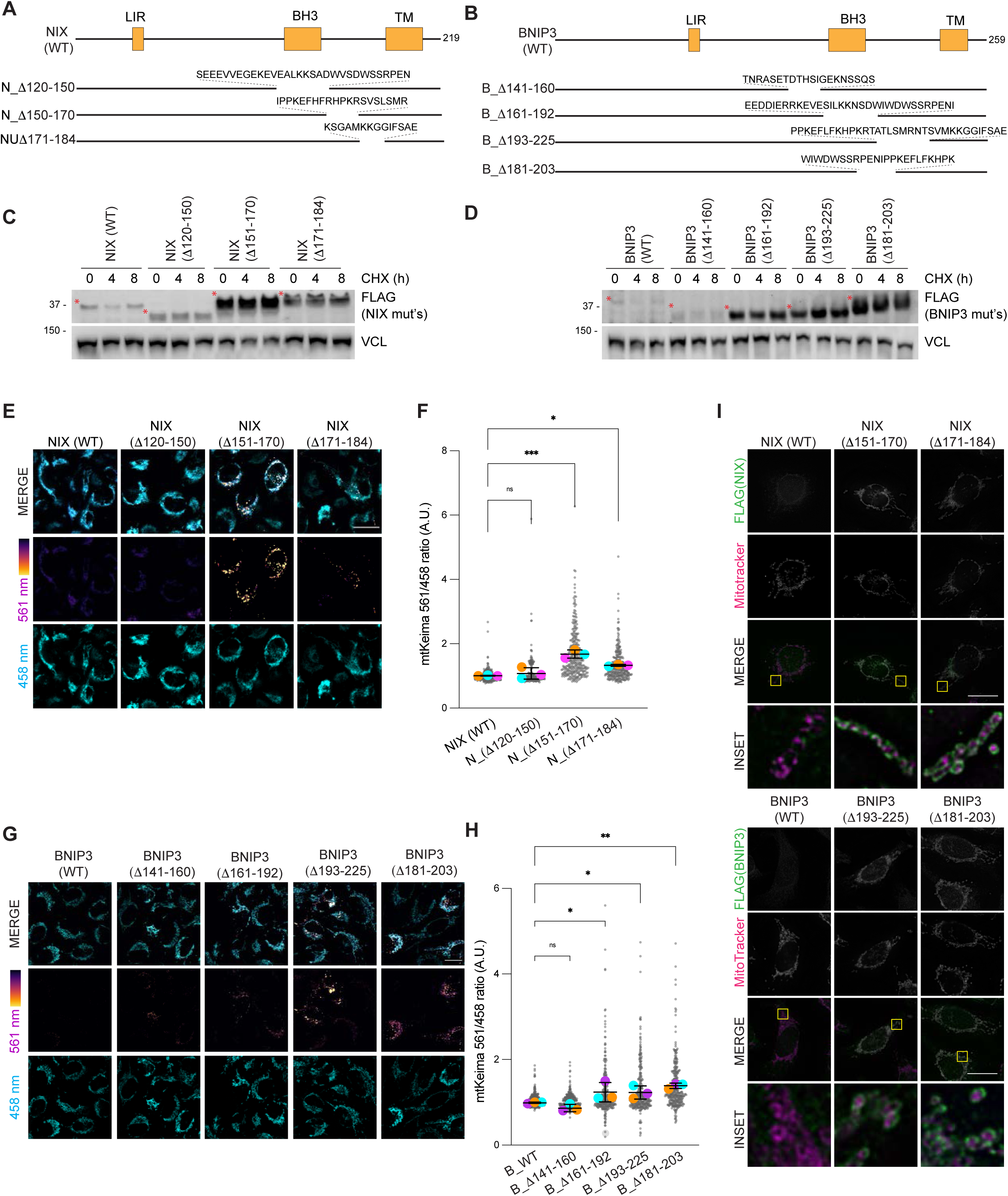
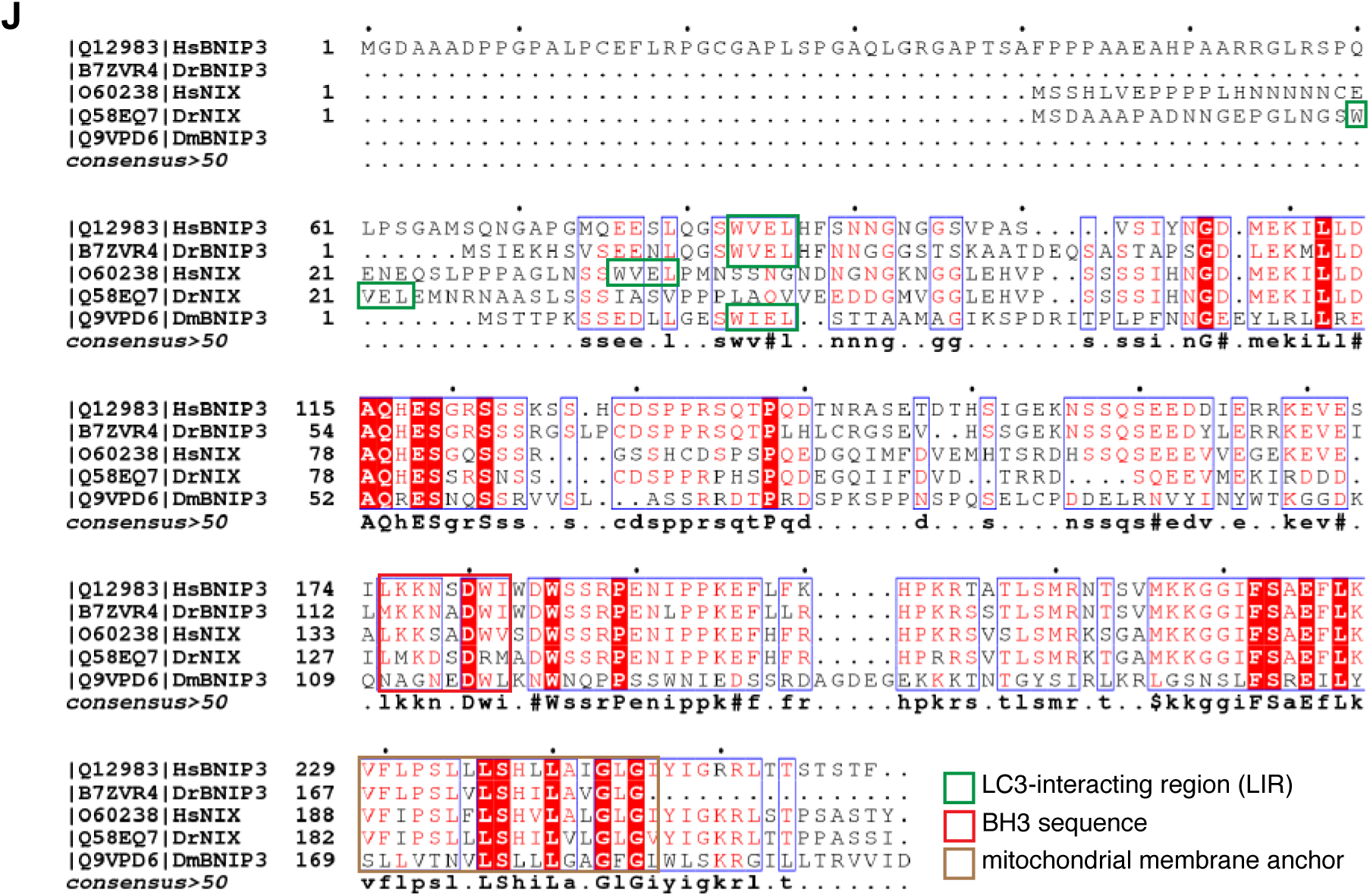
NIX and BNIP3 stabilization promotes mitophagy. A) Schematic representation of full-length NIX and its deletion mutations. B) Schematic representation of full-length BNIP3 and its deletion mutations. C) *C-terminal deletion fragments in NIX have increased steady state and stability compared with wild-type NIX.* Hela-Flp-In NIX knockout cells expressing inducible FLAG-NIX-WT, FLAG-NIXΔ120-150, FLAG-NIXΔ151-170, and FLAG-NIXΔ171-184 were treated with cycloheximide (CHX) for the indicated time. Cells were then lyzed and analysed by immunoblotting. Red asterisks denote the size of NIX or its deletion mutants. D) *C-terminal deletion fragments in BNIP3 have increased steady state and stability compared with wild-type BNIP3.* HeLa Flp-in BNIP3 knockout cells expressing inducible FLAG-BNIP3-WT, FLAG-BNIP3Δ160-183, FLAG-BNIP3Δ181-203, BNIP3Δ201-225 were treated with CHX. Red asterisks denote the size of BNIP3 or its deletion mutants. E) *Inducible expression of hyperstable NIX mutants increases mitophagy.* Hela Flp-in Keima cells stably expressing inducible NIX or deletion mutants were treated with doxycycline for 48 h and mitophagy was evaluated using live-cell confocal fluorescence microscopy. The emission signal obtained after excitation with the 458 nm laser (neutral pH) or 561 nm laser (acidic pH) is shown in cyan or mpl inferno, respectively. F) Quantification of E. N=3. G) *Inducible expression of hyperstable BNIP3 mutants increases mitophagy.* Hela Flp-in BNIP3/NIX DKO Keima cells stably expressing BNIP3 deletion mutants were treated with doxycycline for 48 h and mitophagy was evaluated using live-cell confocal fluorescence microscopy. H) Quantification of G. N=3. I) *FLAG-BNIP3/NIX and deletion mutants localize to the mitochondria.* HeLa-Flp-in cells expressing inducible FLAG-NIX/BNIP3-wild-type or deletion mutants were stained for MitoTracker (red) and FLAG antibodies (green). J) *Alignment of NIX and BNIP3 orthologues outlining conserved residues.* The LC3-interacting region, non-canonical BH3 region, and the trans-membrane domain are shown. The blue boxes represent regions of conservation. White letters on a red background are strictly conserved, and red letters are highly conserved. For data in F and H, mitophagy is represented as the ratio of mt-Keima 561 nm fluorescence intensity divided by mt-Keima 458 nm fluorescence intensity for individual cells normalised to the wild-type condition. Translucent grey dots represent measurements from individual cells. Colored circles represent the mean ratio from independent experiments with > 50 cell analyzed per condition per replicate. The centre lines and bars represent the mean of the independent replicate means +/- standard deviation. P values were calculated using a one-way ANOVA. (\**P*<0.05, \*\**P*<0.005, \*\*\**P*<0.001). ns = not significant. Scale bars = 20 μm.

**Figure EV4.**
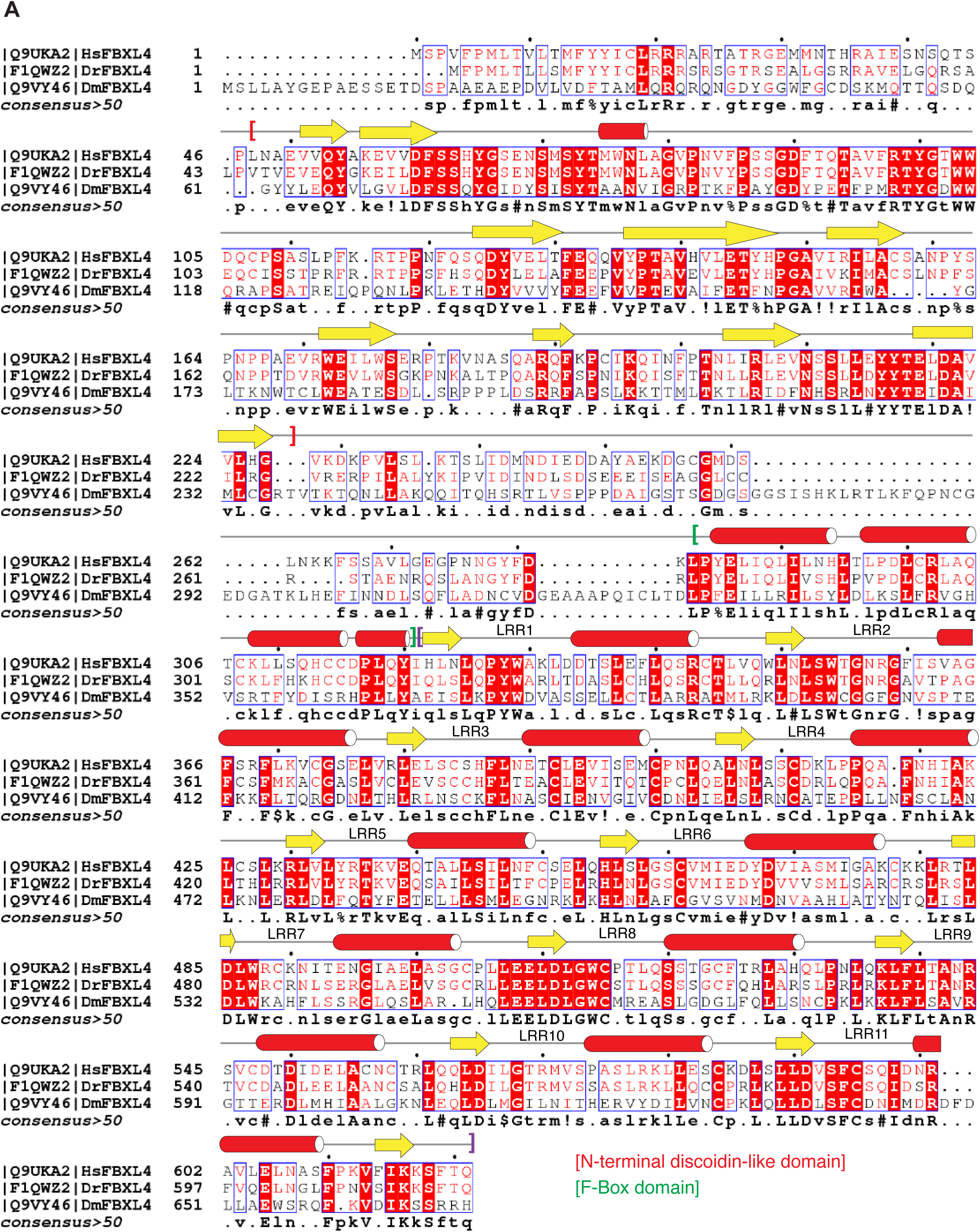
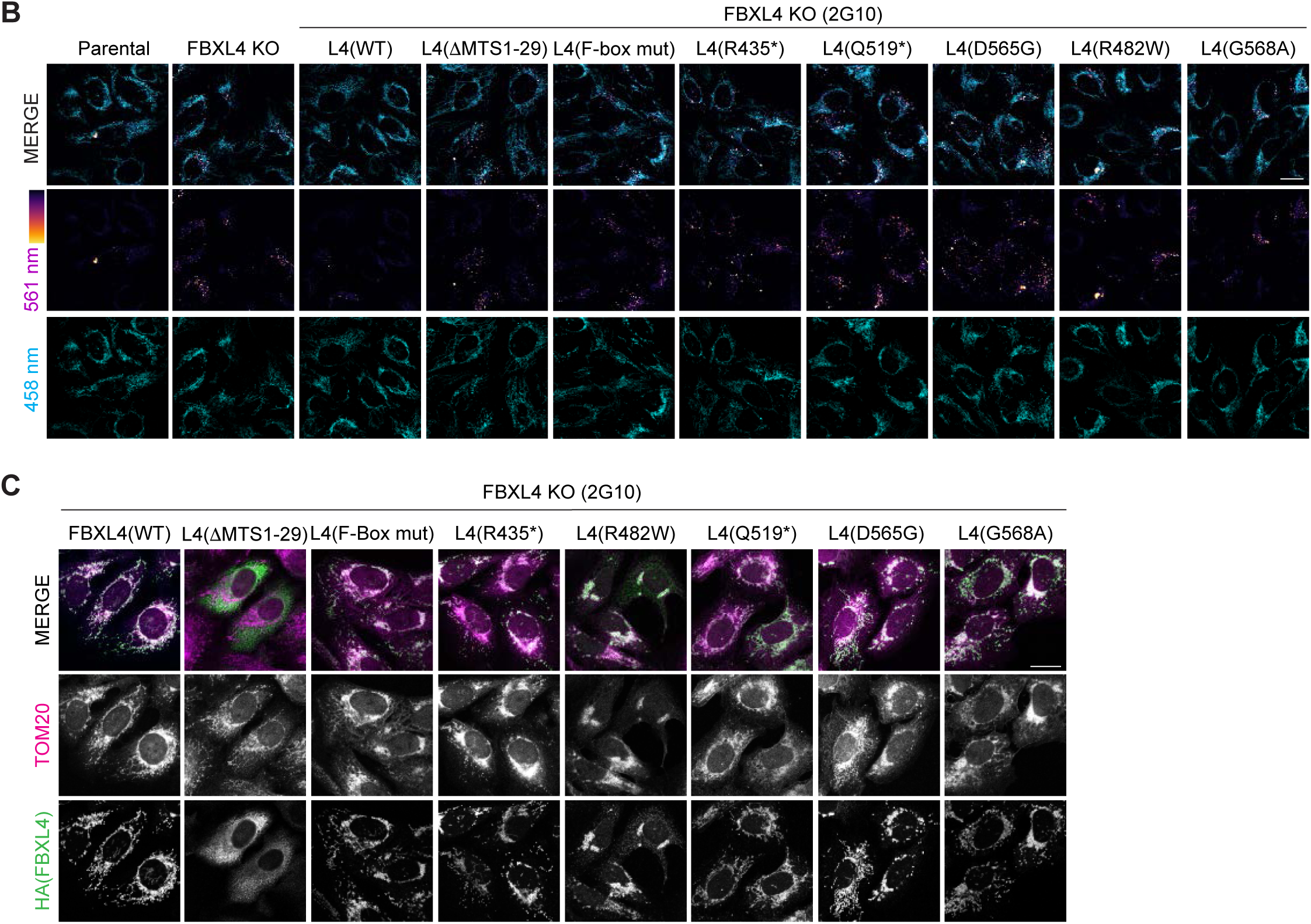
MTDPS13 patient-derived FBXL4 variants do not efficiently assemble into an SCF complex have impaired abilities to mediate NIX and BNIP3 turnover. A) *Alignment of FBXL4 orthologues outlining conserved residues*. B) *FBXL4 patient variants are less efficient than FBXL4-wild-type at suppressing mitophagy.* U2OS mt-Keima cells, U2OS mt-Keima FBXL4 KO 2G10 cells and U2OS mt-Keima FBXL4 KO 2G10 cells rescued with FBXL4 constructs were analyzed by confocal microscopy. The emission signal obtained after excitation with the 458 nm laser (neutral pH) or 561 nm laser (acidic pH) is shown in cyan or mpl inferno, respectively. Quantification is shown in Figure 4C. C) *Localization of FBXL4 variants.* FBXL4 KO cells expressing FBXL4-HA wildtype or specified variants were fixed and stained for HA (to detect FBXL4 in green) or TOM20 (magenta). Scale bars = 20 μm.

**Table EV1.**
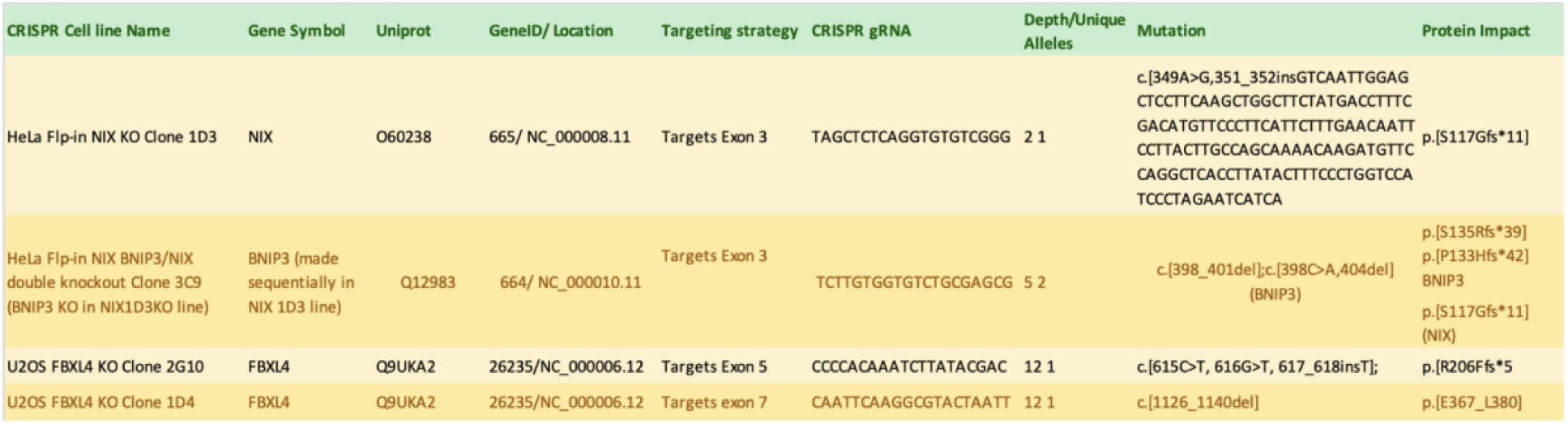
Description of indels detected in the specific CRISPR-Cas9 generated knockout cell lines. Indel mutations and their corresponding mutated proteins (protein impact column) are formatted according to Human Genome Variation Society (http://varnomen.hgvs.org/). The numbers after the asterisks represent the number of amino acids made from the first amino acid changed to the first stop codon encountered.

